# Multiple timescales in bacterial growth homeostasis

**DOI:** 10.1101/2021.03.30.437502

**Authors:** Alejandro Stawsky, Harsh Vashistha, Hanna Salman, Naama Brenner

## Abstract

In balanced exponential growth, bacterial cells maintain the stability of multiple properties simultaneously: cell size, growth rate, cycle time and more. These are not independent but strongly coupled variables; it is not *a-priori* clear which are under direct regulation and which are stabilized as a by-product of interactions. Here, we address this problem by separating different timescales in bacterial single-cell dynamics. Disentangling homeostatic set-points from fluctuations around them, we find that some properties have flexible set-points that highly sensitive to environment - defining “sloppy” variables, while other set-points are buffered and held tightly controlled - “stiff” variables. These control variables are combinations of sloppy ones that compensate one another over long times, creating a hierarchical buffering that protects them from environmental perturbations. This is manifested geometrically as a control manifold in the space of growth and division variables, whose in-plane directions span sloppy variables, while out-of-plane deviations are highly constrained. Cell size is found to be a sloppy variable, which is coupled to growth and division only on the short, single-cycle timescale. Our results show that cellular homeostasis involves multi-level regulation operating on multiple timescales. More generally, our work offers a data-driven approach for identifying control variables in a multi-dimensional system that can be applicable also in other contexts.

## Introduction

Microbial cells are variable in every property that can be measured: size, shape, protein content, metabolic fluxes and more. A clonal cell population is a complex stochastic and nonlinear dynamical system in which cells grow and divide, perpetuating their properties on to the next generation; they interact with their environment as well as with nearby cells. Nevertheless, despite this complexity and variability, a microbial cell population can maintain balanced growth for extended times. In balanced growth, homeostasis keeps the distributions of many phenotypic variables, such as cell size, cycle time and protein content, stable over time – defining a steady state of the system.

Bacterial homeostasis has been the focus of significant research efforts for decades. How is cell size regulated to maintain its stable distribution? How is growth coordinated with cell-cycle events, such as division? These and related questions are central in bacterial physiology; but similar questions arise also in other cases where a large number of interacting variables make up a stable system that maintains homeostasis. The current work reveals that bacterial homeostasis mechanisms span multiple timescales and tie together variables that display a hierarchy of sensitivities to the environment, suggesting principles that might be general to such systems.

Recent years have seen renewed interest in the classic problem of bacterial growth homeostasis (Jun et al. 2018; Meunier, Cornet, and Campos 2021). Following the development of single-cell measurement techniques (Rosenthal et al. 2017), in particular microfluidic single-cell trapping devices (Wu and Dekker 2016) that utilize perfusion to maintain a stable chemical environment, quantitative data could be obtained on different bacterial properties over extended time periods. These data allow mapping distributions of phenotypic variables and correlations among them with unprecedented precision (Campos et al. 2014; Taheri-Araghi et al. 2015; Brenner et al. 2015a; Tanouchi et al. 2015; Grilli et al. 2018). Correlation patterns in these data were studied extensively, and have been interpreted in the framework of stochastic models. Such models assume that cells homeostatically regulate certain properties, e.g. cell size or growth-rate; this allows for comparing and assessing different models for regulation strategy (Osella, Nugent, and Lagomarsino 2014; Amir 2014; Sauls, Li, and Jun 2016; Nordholt, Heerden, and Bruggeman 2020).

In a bacterial cell, many variables are constrained by homeostasis: cell size, protein content, growth-rate per cycle, generation time – all maintain stable distributions over extended times in balanced growth. Importantly, their strong coupling with one another might obscure which variables are actually under regulation, and which are constrained as a result of interactions. An unbiased approach to test this question is required. Recent progress in this direction, using multivariate linear regression, revealed that multiple coordinated mechanisms could be involved in cell size regulation (Kohram et al. 2021). Still, a holistic data-driven approach is not well developed. In particular, homeostasis mechanisms are often assumed to operate within a single cell-cycle time, and correlations are thus examined at the timescale of a single cell-cycle; for example, correlations between initial and final size in a cycle, or added size over a single cycle and its time duration. It is nevertheless possible that homeostasis mechanisms act also on longer timescales. In support of this possibility, recent work revealed that bacterial cellular properties exhibit memory patterns that can extend up to ~ 10 generations (Susman et al. 2018; Vashistha, Kohram, and Salman 2021). Therefore long-term correlations between the different phenotypic variables should also be examined to identify possible slow homeostasis mechanisms.

In this work we characterize bacterial growth homeostasis by studying its sensitivity to perturbations, both external and internal, and by examining correlations among phenotypic variables at different timescales. To this end we distinguish between a homeostatic set-point, empirically estimated as a time-averaged quantity along an individual lineage, and temporal fluctuations around this set-point. The rationale behind our approach is that tightly controlled variables are expected to exhibit robust set-points that are insensitive to perturbations. In contrast, less controlled variables – ones which may be stable due to interactions, but under less stringent direct regulation – may still have homeostatic set-points, but those will be more sensitive and vary with perturbations. This approach is inspired by the concept of robustness in engineering theory, that was widely applied to biology in the context of metabolic and biochemical regulation systems (Savageau 1971; Fell 1992; Stelling et al. 2004). To apply a similar approach to bacterial growth and division, we develop an analysis framework that conditions experimental data and decomposes the ensemble variance, to quantify the robustness of set-points for an array of cellular variables. Because of the large number of parameters the environment depends on (e.g. all medium components, temperature, pH, trap geometry), one cannot easily quantify the perturbations in a set of experiments. Instead, we utilize uncontrolled variation across measurements performed under nominally identical conditions, to reveal the relative sensitivity of the different cellular variables. We analyze data from the recently developed “sisters machine” microfluidic device (Vashistha, Kohram, and Salman 2021); this unique experiment design allows us to distinguish effects of lineage history from micro-environment and to quantify their influence on the set-point variation.

Our results reveal that homeostatic set-points generally vary among individual lineages. Tracing the origin of this variation, we find that the trap environment is a crucial driver of this variation; perhaps surprisingly, the internal cell-state that depends on lineage history, contributes only a small fraction. Despite the uncontrolled nature of environmental perturbations, a qualitatively consistent behavior of cellular variables emerges: the different variables span a broad range of sensitivities to the environment, with a well-defined repeatable hierarchy among them. This hierarchy of sensitivities portrays a multi-variable connected system, forming a cascade of coupling levels that protects some variables from perturbations more than others, a property reminiscent of a sloppy system (Daniels et al. 2008; Braun 2020): in such a system, some variables can vary over a large range (“sloppy” variables), whereas others must be kept tightly controlled (“stiff”variables), to maintain functionality.

Importantly, we find that some sloppy variables co-vary with one another in a simple and logic way over long timescales. Such correlations indicate the existence of long-term hoemostatic mechanisms that extend over multiple cell-cycle times. In the space of growth-rate, generation time and division ratio, (*α*, *τ*, *f*), we identify a control variable hierarchically composed of the sloppier variables, and predict that homeostatic set-points will adhere to a manifold while occupying a large region inside the manifold. This prediction is verified in the entire set of experiments analyzed. These results suggest that long-term growth and division homeostasis is characterized by a “set-manifold” rather than a set-point. Directions on the surface of this manifold are flexible to change with perturbations, while the direction perpendicular to it is stiff. Cell-size is found to be also a sloppy variable, but does not co-vary with any of the growth and division variables over long times. Well-known correlations reported previously are recovered on a cycle-by-cycle basis, and their significance in connection to long-term homeostasis is discussed.

## Results

### Sloppy and stiff variables in trapped bacteria

Despite our best efforts to control the environment in microfluidic devices, some measurements do not exhibit similar statistics in repeated experiments that are nominally identical in external conditions. For example, distributions of cell size and highly expressed proteins collapse onto a universal shape after scaling, over a broad range and with very high precision, but when measured in physical units these variables exhibit a persistent individuality among lineages measured in different traps (Salman et al. 2012; Brenner et al. 2015a; Susman et al. 2018; Kohram et al. 2021). This individuality is apparent well beyond experimental error, but its origins remain unclear. It suggests that individual lineages can maintain stable balanced growth around distinct set-points. To test this idea, we develop a phenomenological approach that quantifies and identifies the source of variation in homeostatic set-points. We apply this approach to different cellular phenotypic variables that can be estimates from cell-size measurements.

Consider individual lineages trapped separately in microfluidic channels – a mother-machine device. Pooling all data from experiments performed under the same controlled conditions makes up the “pooled ensemble”, wherein each measured variable *z* has a total variance *σ*^2^(*z*). This variance is composed of both temporal fluctuations around each trap-specific set-point, and variations of the set-point itself among traps. These contributions can be separated by using the law of total variance:

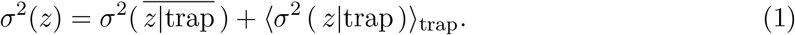

Here 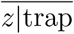 is the average conditioned on the trap, and ⟨·⟩_trap_ denotes the averaging over all traps. The first term is the contribution of different trap averages whereas the second is the contribution of local fluctuations around them, to the total variance over the pooled ensemble. A similar decomposition of variance was previously developed to disentangle contributions to gene expression variation (Bowsher and Swain 2012).

Since mother machine measurements in each trap correspond to a trajectory over time, conditioning on trap is equivalent to a decomposition into temporal average and fluctuations around it: 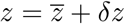. The total variance decomposition then amounts to

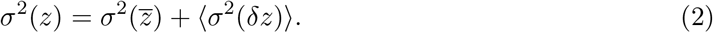

Lineages are statistically equivalent if their temporal averages are all the same and the first term in Eqs. (1), (2) vanishes (or, for a finite data-set, is negligible relative to the total variance). This would be the case if differences between trap conditions and lineage history induce small changes in cell physiology relative to random temporal fluctuations. If we view single lineages as realizations of a dynamical system, then with borrowed terminology, this term reflects weak ergodicity breaking. Indeed, normalized by the total variance, it has been suggested as an ergodicity breaking parameter (He et al. 2008; Meroz and Sokolov 2015), quantifying what fraction of the variance comes from differences among time-averages. In our context we define the trap-conditioned variance fraction

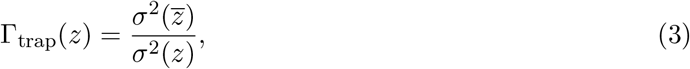

a parameter in the range [0, 1]. Fig. 1 presents an artificial illustration of three groups of trapped bacteria, each group with the same overall (pooled) mean and variance, but different Γ_trap_. For Γ_trap_ = 0 (Fig. 1A), all traces have the same average, and variance is contributed only from fluctuations around this average (the “ergodic” limit). Time series (top panel) mix with one another through the measurements. At the other extreme for Γ_trap_ ≈ 1 (Fig. 1C), variance comes mainly from distinct averages among traps and very little from fluctuations around them. Time series are then seen to center around different means.

**Figure 1:**
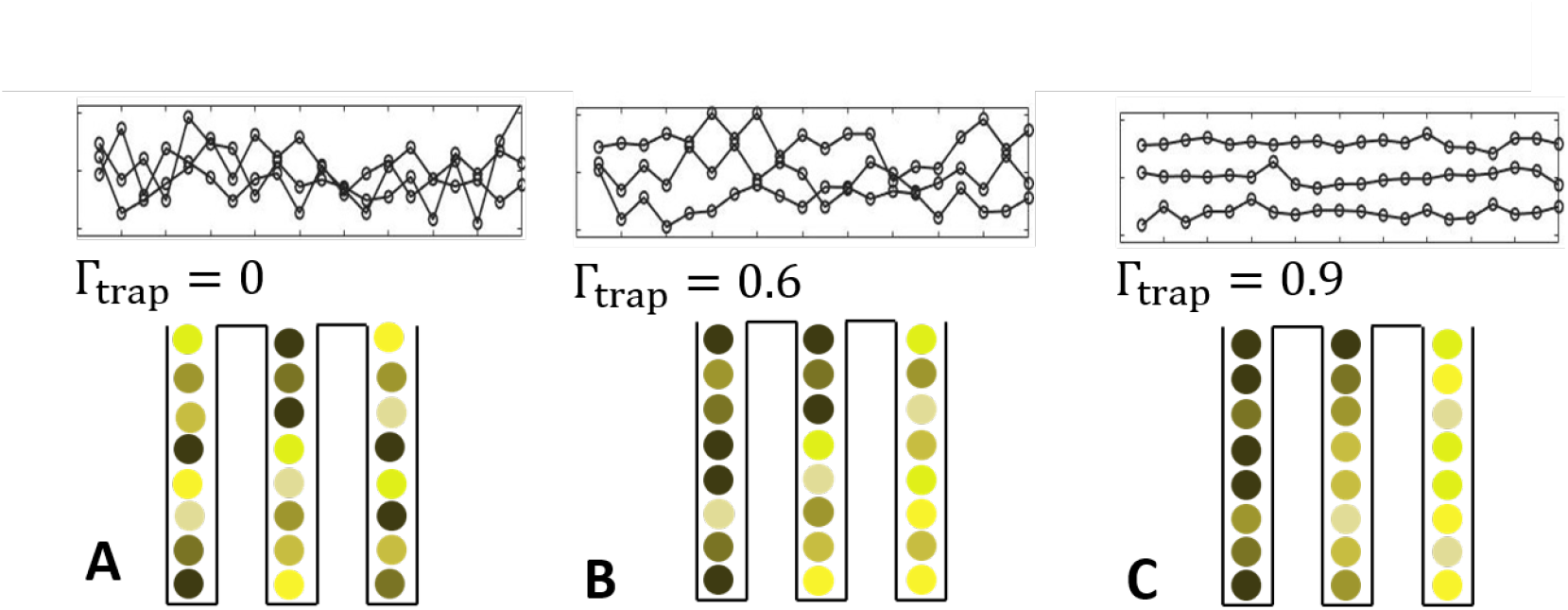
Γ_trap_ quantifies trap contribution to total variance (illustration). Each of the artificial experiments (A,B,C) has zero mean and unit variance when their 3 traps are pooled together. Top: time traces in ensembles with different values of Γ_trap_. It can be seen that as Γ_trap_ increases, distinct trap averages contribute more to the total variance relative to within-trap fluctuations. Bottom: pictorial illustration highlighting the segregation between traps that increases with Γ_trap_.

We apply the variance conditioning to cellular phenotypic variables that can be directly estimated from cell-size measurements over lineages. Such variables have been extensively used to infer statistical properties of bacterial physiology and variability (Amir 2014; Sauls, Li, and Jun 2016; Grilli et al. 2017); they are illustrated in Fig 2A. One may identify the two fundamental time/rate variables in each cycle, the inter-division (or generation) time *τ* and the single-cycle exponential growth-rate *α*. Although growth deviates from an exact exponential when measured carefully (Kohram et al. 2021; Nordholt, Heerden, and Bruggeman 2020; Vashistha, Kohram, and Salman 2021), the best fit exponential rate is still a useful approximation for growth over the cell cycle and will be used in what follows. Cell size is characterized by its length *x*_0_ at the start and *x*_*τ*_ at the end of each cycle, as well as their difference Δ = *x*_*τ*_ – *x*_0_.

**Figure 2:**
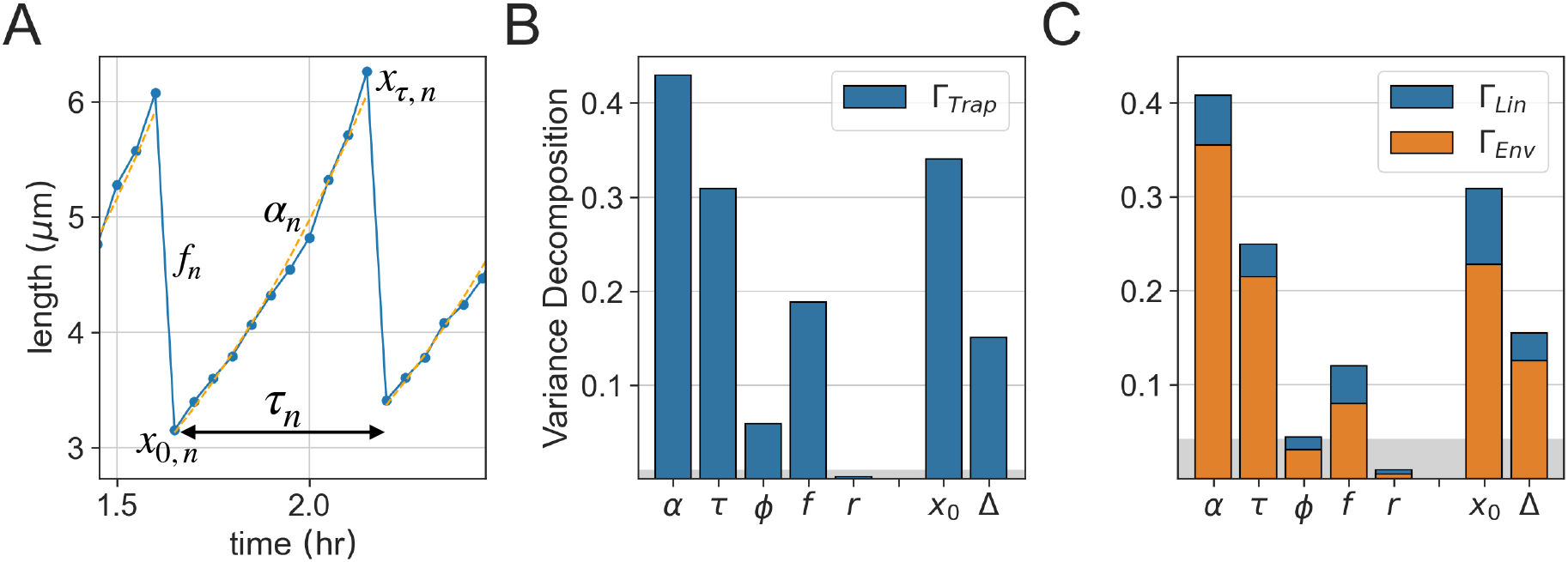
Distinct set-points (time-averages) contribute to total variance. **(A)** Illustration of the main phenotypic variables in a cell cycle. Presented for the *n*-th cycle are the initial size (*x*_0_, *_n_*), size at division (*x*_*τ*_, _*n*_), exponential growth-rate (*α_n_*), inter-division time (*τ_n_*) and division ratio 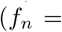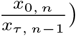. Aditional variables are the total growth (*τ*_*n*_ = *α*_*n*_ *τ*_*n*_), added length (Δ_*n*_ = *x*_*τ,n*_ – *x*_0, *n*_) and size ratio 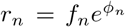. **(B)** Variance decomposition conditioned on trap for a set of mother machine experiments performed under identical conditions. Bar heights are the fraction of variance coming from differences among trap-specific set-points. **(C)** Variance decomposition conditioned on trap micro-environment (orange) and on lineage identity (blue), computed from a set of sisters machine experiments under identical conditions. In both **B** and **C**, the remaining fraction complementing to 1 represents the contribution of temporal fluctuations to the normalized variance. The grey shadow is the noise level (finite size sampling effect) estimated by running the same analysis on artificial lineages drawn at random from the pooled ensemble (See Methods).

Finally, several relative dimensionless size variables can be defined: the logarithmic fold-growth per cycle *ϕ* = *ατ* = ln(*x*_*τ*_/*x*_0_); the fold-decrease at division *f*_*n*_ = *x*_0_(*n*)/*x*_*τ*_(*n*–1); and the total relative change in cell size over an entire cycle 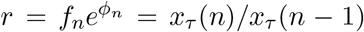. The last two connect consecutive cell cycles, cycle *n* – 1 to cycle *n*. We note that this is not a set of independent variables; on the contrary, some are clearly combinations of others. We investigate them separately and in relation to one another, with the aim of obtaining an unbiased phenomenology that will reveal their significance in the cellular context. The global CV (standard deviation divided by mean) of all these variables over pooled ensembles are listed in SI Table 2.

Using the variance conditioning described above, we estimated Γ_trap_ for each of the phenotypic variables in a set of pooled mother machine measurements; they are plotted as bars in Fig 2B. The remaining fraction (complementing the bar heights to 1) represents the contribution of temporal uctuations around individual trap-conditioned time average. Finite-size sampling effects, estimated from a set of artificial lineages with the same global statistical properties, is depicted as a control (gray baseline; see Methods).

Growth-rate *α* shows the largest trap-dependent variance, with more than 0.4 of its total variance contributed from differences in temporal means among traces. Generation time *τ* is also significant with Γ_trap_ ≈ 0.3. Interestingly, their product - fold growth *ϕ* - shows a markedly smaller value of Γ_trap_, indicating that its average is more strongly regulated across traps. Similarly, the initial size *x*_0_ shows a larger value than Δ, as does *x*_*τ*_ (not shown). Finally, the product *r* = *fe*^*ϕ*^ also shows a smaller value of Γ_trap_ than each one of its factors, and moreover, even smaller than the noise level. These observations hint to nontrivial correlations of variables across time in each trace, as will be discussed in detail below.

### Sensitivity to environment underlies sloppy set-points

Before addressing the relationships between variables, we first ask: What is the source of the variability among individual trap averages? Possible sources can be broadly categorized into two types: internal and environmental. Internal individuality reflects the lineage history, epigenetic memory, and any hidden internal cellular variables that are at least partly inherited across divisions (for example, age of the mother cell at the beginning of the measurement). Environmental factors include variation in physical variables such as channel width and temperature, as well as local biochemical variation in growth medium. Mother machine data cannot distinguish between these two sources.

We utilize the experimental design of the sisters machine to follow two lineages in each channel of a single trap, which start growing at roughly the same time (Vashistha, Kohram, and Salman 2021). In this device, lineages stemming from two sister cells – “sister lineages” – were previously used to analyze the divergence in the physiological variables between two sisters after their separation (Vashistha, Kohram, and Salman 2021). Here, we examine pairs of unrelated lineages having flown into both sides of the same trap and grown in parallel in that trap – “neighbor lineages”. Conditioning of the total variance on trap now includes two separate lineages with different histories but a shared environment. Refining our variance decomposition to include these two sources, we obtain three components:

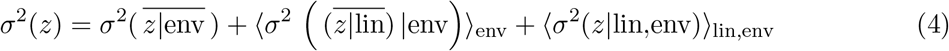

(See Methods for details). Here, the three terms describe the contribution to total variance of trap environment, individual lineages given the environment, and fluctuations within individual lineages around their average. Normalizing by the total variance we find that Γ_trap_ splits into two terms, Γ_trap_ = Γ_env_ + Γ_lin_.

Fig. 2C depicts this decomposition with the two fractions Γ_env_, Γ_lin_ marked respectively by orange and blue. As before, the remaining fraction complementing to one is the contribution of temporal fluctuations around individual lineage averages. For most variables, the contribution of lineage individuality is approximately 2 – 5% (blue part of bars). With the exception of *r*, the trap environment is the major source of conditioned variance to all variables in the set of experiments analyzed (orange portion of bars).

Comparison of Figs. 2B and 2C provides an important test for the generality of our results: disregarding the internal decomposition into two sources (two colors) in C, the total bar heights, representing Γ_trap_, should be similar. Recall that the data-sets are taken from two different microfluidic machines; while nominal conditions are the same, in each pooled ensemble the collection of traces is characterized by an uncontrolled variability in environments - everything that is beyond the experimenter’s control (for further dependence on experiments and the effect of pooling see SI Figs. 8–9). It is not surprising therefore that the results are not identical. However, a considerable level of repeatability is seen in the *relative* behavior of the different phenotypic variables. It appears that some are regulated more tightly around a well conserved set-point, which is robust to both environmental and internal variability; most pronounced are the variables *ϕ* and *r*. In contrast, growth-rate, inter-division time and size variables have a significant fraction of their variance contributed from different time averages, suggesting they are sensitive to perturbations and regulated only locally around different set-points. For further comparison, SI Figs. 9,10 show the same analysis for previously published results covering two bacterial strains (Wang et al. 2010) and a range of temperatures (Tanouchi et al. 2015).

We next return to the correlations that arise among set-points along time, and their implications to growth and division homeostasis.

### Long-term homeostatic correlations in growth and division

The large contribution of micro-environment to the variance of growth-rate *α*, seen in Fig. 2B,C, indicates that it fluctuates around distinct set-points in different traps. A direct way to visualize this environment dependence is to examine the time averaged growth-rate of two neighbor-lineages grown in the same trap in the sisters machine; Fig. 3A shows that they are indeed correlated. The diagonal spread of points represents the range across traps, and is larger than the off-diagonal spread representing variation between one lineage and its neighbor in the same trap. Neighbor cells are correlated only in their set-points and not in the fast fluctuations around them (SI Fig. 15), indicating that the effect of the environment has a long timescale closer to the lineage length than to the single cell cycle time. A qualitatively similar picture emerges for inter-division time, as shown in Fig. 3B.

**Figure 3:**
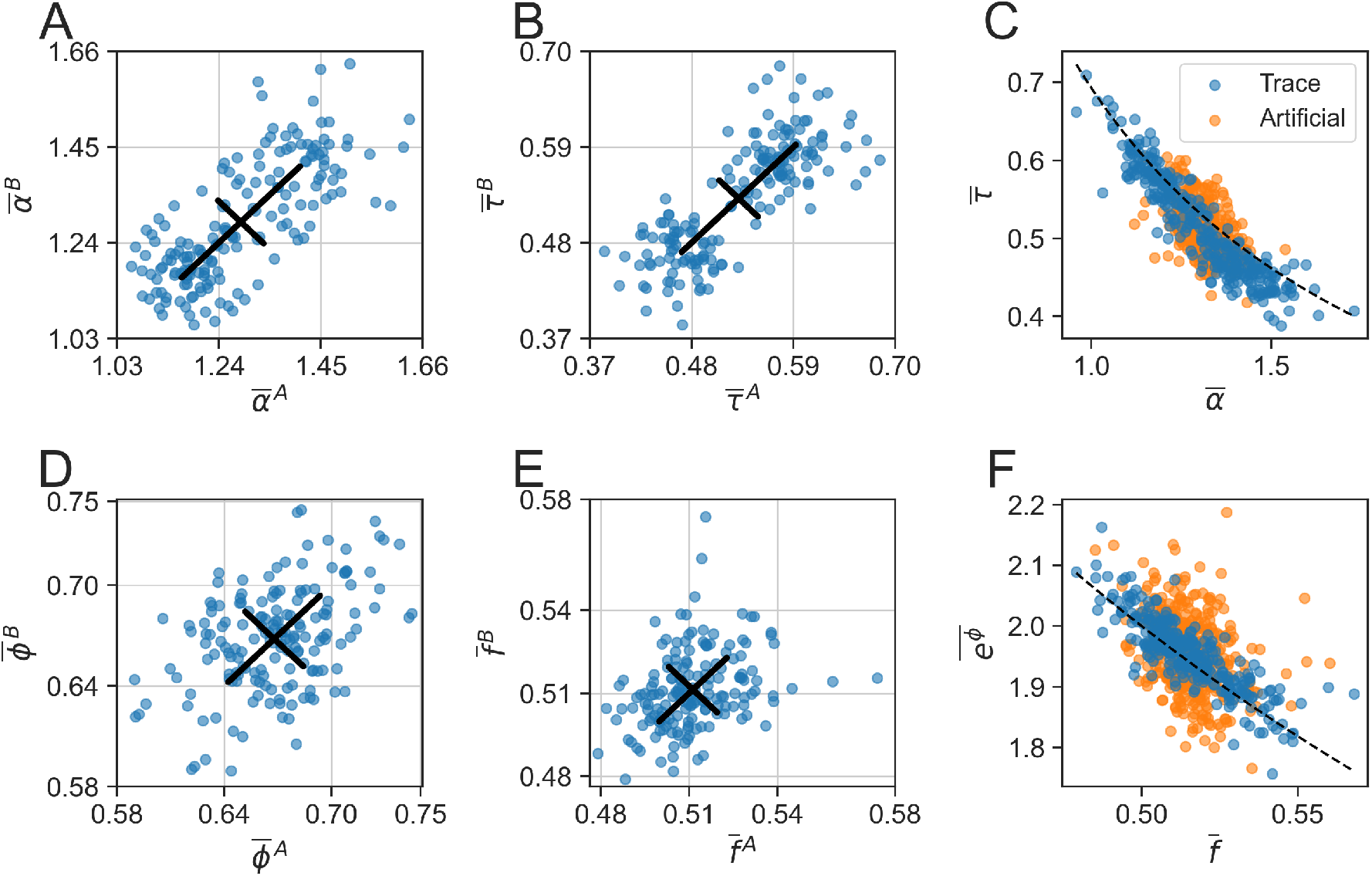
Co-variation of sloppy homeostatic set-points. (**A,B**) Sloppy variables *α, τ* display a range of homeostatic set-points (time averages). Scatter-plots for two neighboring lineages in the same microfluidic trap: **(A)** Exponential growth-rate *α*, Pearson *ρ* = 0.74; **(B)** generation time *τ*, *ρ* = 0.8. The set-points are primarily sensitive to the environment (diagonal spread), and less so to lineage history (off-diagonal spread). Black lines are standard deviations in the two directions. **(C)** Within each lineage, set-points strongly co-vary (blue circles; *ρ* = −0.9) and lie close to the line *ατ* = ln 2 (dashed black). Orange circles show averages of per-cycle variables over arbitrary groups <u>o</u>f the same sizes as lineages (artificial lineages; *ρ* = −0.42). **(D)** Scatter-plot of fold<u>-g</u>rowth set-points 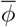 for neighbor cells, Pearson *ρ* = .43. **(E)** Scatter-plot of division fraction set-points 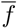 for neighbor cells, Pearson *ρ* = .33. **(F)** Same presentation as in **C**, for the variables 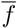 and 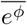. The dashed black line is 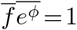; *ρ* = −.84 for temporal averages, *ρ* = −.38 for artificial averages. For a full matrix of set-point scatter-plots of pairs of variables between neighbor lineages, see SI Fig. 14.

By common intuition, homeostasis requires no drift in growth and division across time. With symmetric cell division, this would dictate that cells roughly double their size over the cell-cycle. This, in turn, requires the set-points (time-averages) of growth-rate and inter-division time to be correlated within each lineage, so that 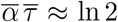; Fig. 3C (blue circles) shows that indeed the set-points are located along this expected line (black dashed line).

Correlations between growth-rate and inter-division time were previously reported over short-times, on a cycle-by-cycle basis (Grilli et al. 2018; Panlilio et al. 2020). Typically such correlations are inferred from a large number of data points, each representing a single cell-cycle from the pooled ensemble, making up a noisy cloud; in our experiments, this set of points has a Pearson correlation coefficient of *ρ* = −0.52. Had the per-cycle correlations been the only mechanisms at play, correlation between averaged values would reflect a merely statistical effect: they would remain qualitatively the same as the per-cycle correlations, with a possible decreased magnitude due to the smaller dynamic range after averaging. This effect is depicted in Fig. 3C by the orange symbols, where arbitrary finite samples from the pooled ensemble were averaged (artificial lineage averages; *ρ* = −0.42). In contrast, averaging over time in single lineages, we find that the correlation coefficient is dramatically increased, reflecting the large range of distinct and correlated time-averages (blue circles; *ρ* = −0.9).

These differences between the pooled and time-averaged correlations suggest a long-term mechanism that is obscured by per-cycle noise and smoothed when looking at time-averaged variables. One may assign quantitative measures to the contribution of short vs. long term mechanisms by decomposing the covariance cov(*α*, *τ*), similar to the variance decomposition employed earlier. To obtain a dimensionless measure we normalize the covariance by the pooled-ensemble standard deviations; we find this normalized covariance of −0.52 decomposes into −0.28 from long-term and −0.24 from short-term processes (see SI Fig. 11). We conclude that *α* and *τ* are negatively correlated on both long (multiple cycles) and short (single cycle) timescales. Note that the normalized measure decomposes the Pearson correlation over the pooled ensemble into two parts, neither of which is a Pearson correlation in itself. The high value of 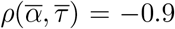 between time averages appears when normalizing correctly by their respective standard deviations 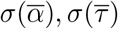 (see Methods).

SI Fig. 11 presents the covariance decomposition for all pairs of variables. This decomposition allows us to disentangle the contributions of long- and short-time processes to the coupling of any two phenotypic variables; we have seen that for *α* and *τ* both contribute, but this is not always the case, as discussed below. We note that despite the numeric difference between components of the covariance and Pearson correlation coefficient, the two measures are mostly qualitatively consistent (SI Fig. 11 compared to Fig. SI 12).

As the set-points of *α*, *τ* lie close to the line of constant product, it is of interest to examine next the product itself: 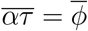 is the logarithmic fold-growth in cell size per cycle. This variable is also affected by the trap environment, but to a lesser extent (Fig. 3D). The range across different micro-environments is now more similar to the range between neighbors in the same trap; the scatter-plot is closer to circular. This is just a different way of expressing the smaller relative contribution of trap environment to conditioned variance of *ϕ* compared to *α* and *τ*, that was seen in Fig. 2A,B. A similar behavior is displayed also by the fraction *f*, shown in Fig. 3E, again reflecting that this variable keeps its set-point more consistent among individual traps.

Applying the no-drift criterion to *ϕ*, *f* suggests that deviations from *ϕ* = ln 2 might be compensated by deviation of exactly symmetric division. Indeed, Fig. 3F shows that set-points of division ratio and fold-growth are correlated and lie along the line 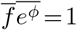. This result is somewhat surprising, as *f* exhibits the smallest CV among phenotypic variables, and is often considered irrelevant to growth homeostasis. A recent study however, showed that on short timescales *f* does correlate with the growth rate in the following cell cycle: the smaller-fraction daughter cell was found to grow faster than its larger sister (Kohram et al. 2021).

Examining the decomposition of cov(*ϕ*, *f*) into short and long timescales (Fig. SI 11), we find that the largest contribution comes from short-term fluctuations around the set-point, approximately 80%. This mostly reflects the small dynamic range of averages 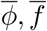, relative to the range of per-cycle fluctuations. After normalizing by their respective standard deviations, we find that the time-averages have a high Pearson correlation coefficient, 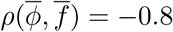.

The product of *f* · *e*^*ϕ*^ = *r* makes up the ratio between cell sizes over a full cycle. Looking back at Fig. 2 we recall that this variable is the tightest of all, holding a strict set-point whose contribution to variance is even smaller than the noise level (Figs. 2B,C). We interpret this result to imply that over long times this variable is tightly regulated to maintain an absolute constant set-point, despite variability of its factors. Taken together, these results point to a two-level hierarchy among phenotypic variables: while the set-points of *α*, *τ* are free to drift with changes in the environment, their product *ϕ* is more constrained; at the next level, this product is coupled to the division fraction variable *f*, and their product is in turn even more strongly buffered from perturbations. These processes take place on long timescales of many generations and are revealed by averaging individual traces over sufficiently long time. Looking in the three dimensional space (*α*, *τ*, *f*), Fig. 4 shows that the lineage set-points adhere to the manifold *r*(*α*, *τ*, *f*) = *f e*^*ατ*^ = 1 (4A, side view of manifold); the spread perpendicular to this manifold represents the variability in set-points of *r*, which is precisely its (negligible) conditioned variance in Figs. 2B,C. In contrast, the variables that make up *r* span a significant range inside this manifold (4B, front view of manifold) . Most points are close to the line *ατ* = ln 2; those that deviate are still on the manifold, with values of *f* removed away from 1/2 to compensate and hold *r* still close to 1.

**Figure 4:**
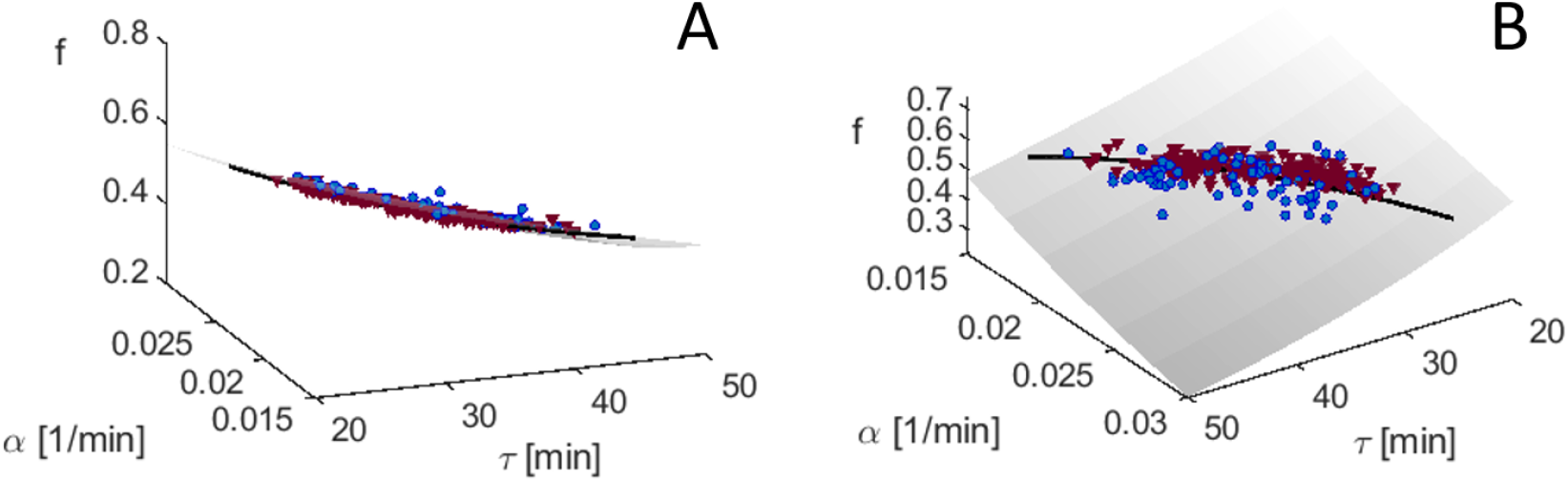
Homeostasis attractor manifold, two views. Homeostatic set-points (temporal averages) of lineages from mother machine experiments (blue circles) and from sisters machine experiments (violet triangles), both grown in LB medium at 32°. **(A)** Side view, emphasizing the small variability in the direction perpendicular to the manifold. **(B)** Front view emphasizing the spread within the manifold, in the vicinity of *ατ* = ln 2 (black line). In both panels, gray manifold represents *r*(*α*, *τ*, *f*) = *fe^ατ^* = 1.

### Cell Size variables

As seen in Fig. 2, size variables *x*_0_ and Δ show a significant conditioned variance, indicating sloppy set-points across trap environments. This extends and quantifies the results of Susman et al. 2018. The different measures of cell size exhibit long-term correlations among themselves, that follow from general intuitive homeostasis arguments. For example, assuming approximately symmetric division, one must have on average 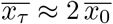. Fig. 5A shows that measured lineage time-averages keep this relation to a good approximation (blue circles, with the black dashed line showing the expected relation). This highlights the stability of cell-size homeostasis along time, concomitant with a broad range of size set-points maintained in individual lineages -from 2 to 3.5*μm*. We compare this to previously studied correlations between per-cycle data over the pooled ensemble, plotted as orange contours in Fig. 5A. Interestingly, these show a different positive relation, with a slope closer to 1: *x*_*τ*_ ≈ *x*_0_ (orange contours). This relation characterizes also the binned per-cycle data (black squares) and their averages over artificial lineages (orange circles), consistent with previous publications, where size progression over a cycle was approximated by a linear map with a slope close to 1 (Amir 2014; Soifer, Robert, and Amir 2016; Tanouchi et al. 2015).

**Figure 5:**
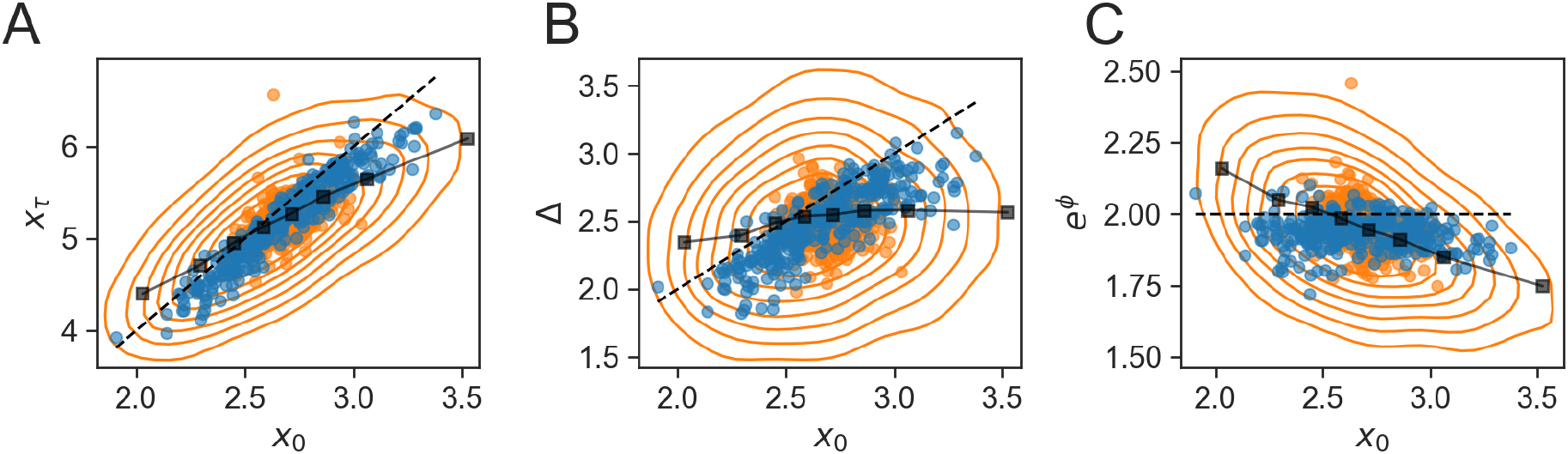
Cell size correlations over long and short timescales. Correlations between initial size and **(A)** size at division, **(B)** added size, and **(C)** fold growth. The orange contours represent percentiles 20 to 80 of the pooled ensemble per-cycle data, black squares are the same data after binning. Blue and orange circles are the averages of real and artificial lineages respectively. Black dashed lines represent expected relations from general considerations of long-term homeostasis: *x_τ_ =*2*x*_0_ **(A)**, Δ = *x*_0_ **(B)**, *e*^*ϕ*^ = *x*_τ_/*x*_0_ = 2. The Pearson correlation coefficients for the pooled ensemble, artificial and lineage set-points respectively are: .61, .64 and .95 for **(A)**, .1, .17 and .82 for **(B)** and −.43, −.35 and −.11 for **(C)**. coefficients for lineage averages, the pooled ensemble,

In terms of the added size Δ = *x*_*τ*_ – *x*_0_, the difference in correlation slopes found in Fig. 5A is now transformed into a qualitatively different behavior of pooled vs. time-averaged data. Fig. 5B shows that the average values follow approximately Δ ≈ *x*_0_ (blue circles with expected black dashed line), consistent with 5A. In contrast, the pooled data shows no correlation between Δ and *x*_0_ – the well-known adder correlation pattern, reported in several types of bacteria (Sauls, Li, and Jun 2016; Soifer, Robert, and Amir 2016; Willis and Huang 2017; Yu et al. 2017; Eun et al. 2018). Adder correlations are seen also in the binned data and in averages over artificial lineages (black squares and orange circles), but not in the time averages (blue circles). We see here our first example of qualitatively different correlations between the same phenotypic variables on the short and long timescales (there will be more below). How do such qualitative differences come about?

Fig. 5C might provide a clue to answer this question. Previous work has shown that the adder correlation – the independence of added volume on initial size – is equivalent to a negative correlation between fold-growth and initial size (for a derivation see Appendix in Susman et al. 2018). Such negative correlation was reported several times in previous work (Amir 2014; Taheri-Araghi et al. 2015; Brenner et al. 2015b; Grilli et al. 2018; Susman et al. 2018). Indeed we observe it in the pooled data (orange contours in Fig. 5C), also after binning (black squares) and averaging over artificial lineages (orange circles). However, when averaging over lineages this correlation is lost; the set-points of *x*_0_ and fold-growth *e*^*ϕ*^ (blue circles), show no significant correlation (*ρ* = −0.11). Examining the figure it can be appreciated that, relative to the range of single-cycle value (contours), 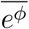 is a stiff variable that spans a narrow range across lineages, while 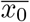 spans almost the entire range. It therefore seems that the lack of correlation over long times is in this case related to the large differences in range of set-points between the sloppy cell-size and the stiff fold-change. The residual variability in the set-point of *e*^*ϕ*^ is not significantly correlated with that of *x*_0_.

More generally, however, set-points of cell-size variables are not significantly correlated with any of the growth variables (*α*, *τ*, *f*) over long timescales, as seen in Fig. 6 (A,B,C blue circles); the correlation is small despite the fact that there is no dramatic difference in the sloppiness of the variables. In fact, long-term homeostasis does not require the connection of size to growth set-points: the intuitive arguments on relations between growth variables invoked above (e.g. *ατ* ≈ ln 2), are indifferent to the cell size itself and only refer to relative sizes. Similarly, homeostatic relations between cell size variables themselves (such as *x*_*τ*_ ≈ 2*x*_0_), do not contain reference to temporal variables or to division. Therefore, correlations between size and growth/division variables represent specific and short-term mechanisms within the cell cycle. Indeed, recent work points to specific processes such as protein accumulation to threshold for completion of discrete cycle events, that underlie such correlations (Si et al. 2020; Nordholt, Heerden, and Bruggeman 2020; Kohram et al. 2021). Taking into account that lineages maintain distinct homeostatic set-points, we expect such mechanisms to operate on *relative* variables - and to be seen most clearly when set-points are subtracted. The bottom row in Fig. 6 illustrates these short-term correlations: for each variable, the lineage-specific set-point is subtracted and correlations are plotted over the pooled ensemble. As expected, we find that compared to the original pooled data (Fig. 6, top row), correlations are enhanced when looking at relative variables. For example, while *ρ*(*x*_0_, *τ*) = −0.3, in relative variables *ρ*(*δx*_0_, *δτ*) = −0.4.

**Figure 6:**
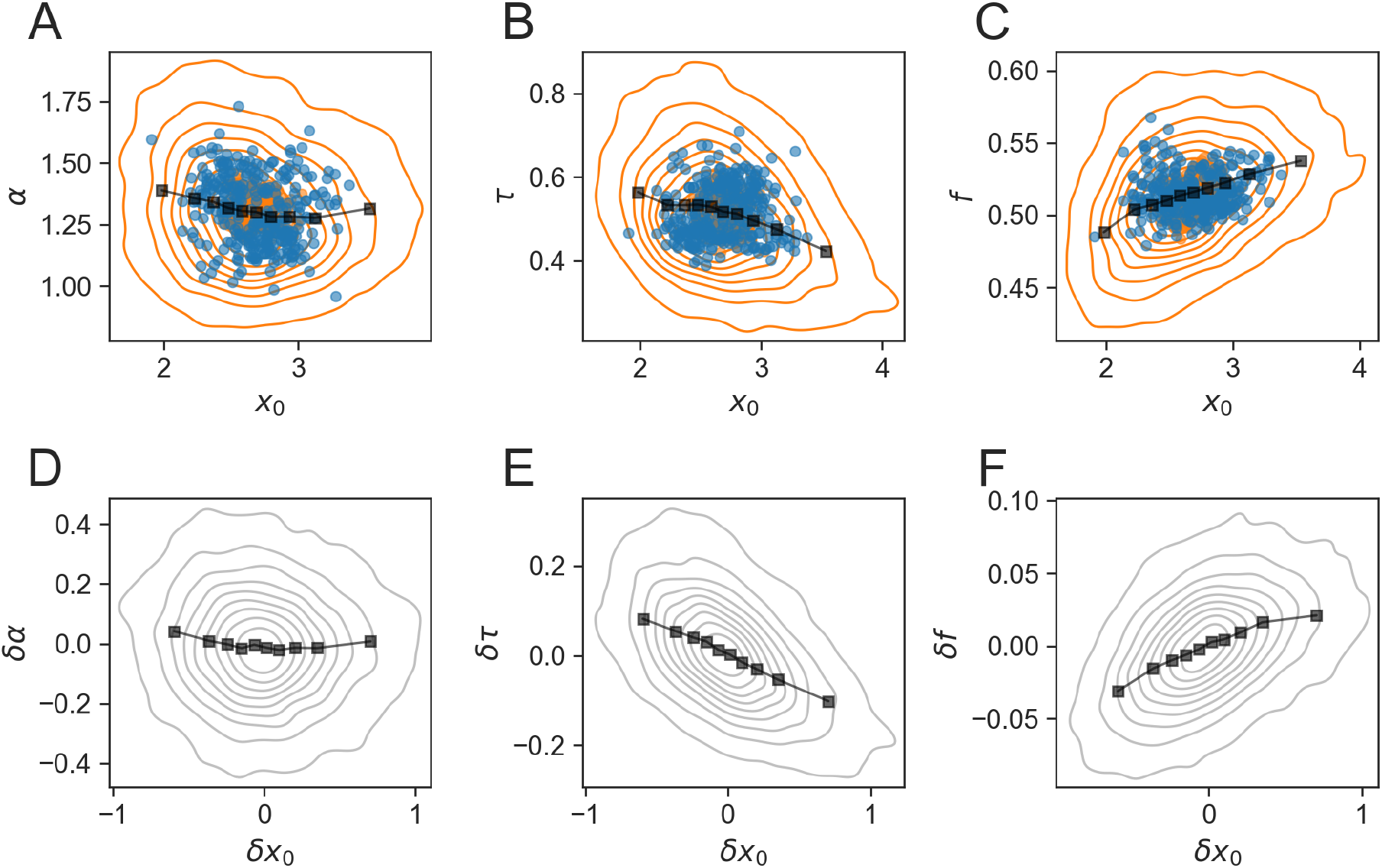
Correlations between cell size and growth/division variables. Top: contour plots of correlation between pooled per-cycle variables in physical units (orange contours; *ρ* = −0.12, −0.3, −0.37 for A,B,C respectively), their binned values (black square) and time averages (blue circles; *ρ* = −0.19, −0.1, −0.11). Bottom: Correlations between relative variables, where lineage set-points (long term averages) have been subtracted, sharpen the top row correlations (*ρ* = −.05, −.49, 0.44 for D,E,F.). Grey contours: pooled data for relative variables, black squares: binned data.

In a similar approach, all pairs of phenotypic variables were divided into their time-averages and relative variables, and the correlations between them examined separately. The results are summarized in SI Figs. 12 and 13. Some correlations stem from a simple mathematical dependence, for example *ρ*(*ϕ*, *τ*) reflects the fact that these variables are proportional to one another; others highlight the separate contributions of long and short timescales in the coupling between different variables.

### Persistent and anti-persistent temporal fluctuations

The previous analysis suggests that short-term processes over the single cell cycle are best seen in relative variables, after subtracting the lineage-dependent time average. Correlations over the ensemble are generally sharpened, as the confounding effect of pooling together variables with different averages is removed. However, the correlations across the ensemble do not tell the whole story – we next focus on the dynamic aspect of the temporal fluctuations.

It is often assumed that if a quantity fluctuates, it accumulates error and can cause instability; this intuition comes from independent fluctuations, and is not necessarily the case in the presence of correlations. We use Detrended Fluctuation Analysis (DFA) to measure the degree to which fluctuations of a variable *δz_n_* accumulate (Peng et al. 1995). Given a time-series of fluctuations in a lineage of length *N*, let 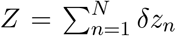 be the cumulative sum which, if the fluctuations were independent, would accumulate like a random walk. DFA estimates the standard deviation of fluctuations *F* (*k*) around linear approximations of *Z* in windows of size *k*, as a function of window size (see Methods). This quantity generally increases with *k* as a power law, *F*(*k*) ∝ *k^γ^*; *γ* ≃ 0.5 corresponds to a sequence of independent variables, 1 > *γ* > 0.5 corresponds to a positive correlation between consecutive steps (a persistent signal), and *γ* < 0.5 corresponds to a negative correlation between consecutive steps (anti-persistent).

Fig. 7 shows the results of this analysis. Fig. 7A depicts the outline of the calculation, whereas 7B shows the scaling of *F*(*k*) with window size *k*. For initial size *x*_0_, the scaling is approximately ~ *k*^0.8^ whereas for *ϕ* the power is smaller than 1/2. For all phenotypic variables considered, the scaling power is plotted in 7C, together with the control estimation of shuffled lineages; all of these show a power of 1/2 as expected from uncorrelated variables.

**Figure 7:**
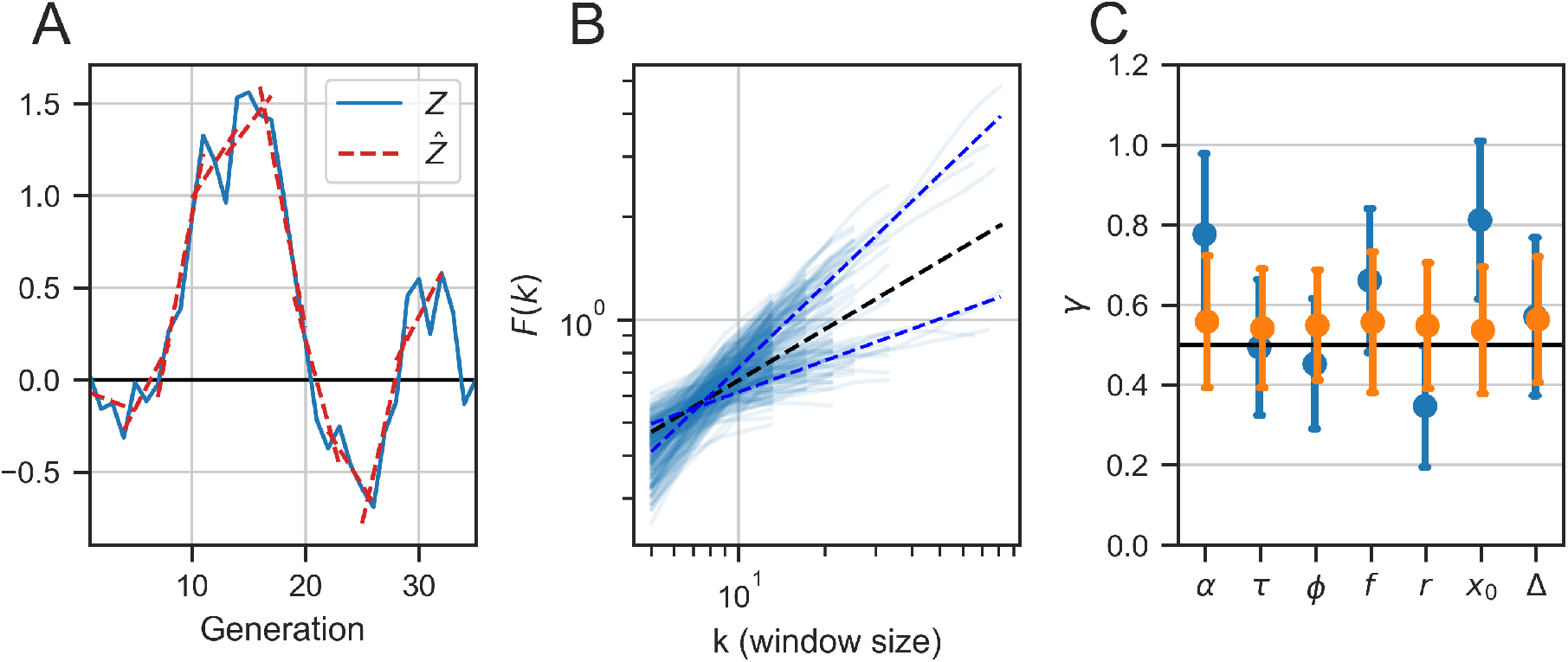
Persistence of phenotypic variables. **(A)** An illustration of the normalized cumulative sum (blue solid line) and its linear approximation for windows of size *k* = 5 (red dotted line). *F* (*k*) will estimate the standard deviation around the linear fits as a function of window size. **(B)** Standard deviation of the fluctuations *F* (*k*) as a function of window size *k* for each lineage separately (light blue lines) and averaged over all lineages (dotted blue lines), showing approximately *F* (*k*) ∝ *k^γ^*. Top line: *x*_0_, with power *γ* = 0.81; bottom line: *r*, with *γ* = 0.3. The black dotted line illustrates *F* (*k*) ∝ *k*^0.5^, which is what is expected from temporally independent fluctuations. **(C)** Mean and the standard deviation of the scaling exponents of each variables in individual lineages from mother and sisters machine lineages (No. 0 – 7 in SI Table 1; blue symbols). Orange symbols show the same analysis for lineages with shuffled generation order. See SI Fig. 17 for the same analysis on publicly available data-sets.

The first conclusion from Fig. 7C is that the phenotypic variables behave differently from one another also in the temporal aspects of their fluctuations around set-points. Moreover, we find that there is an approximate correspondence between sloppiness of set-point and accumulation of fluctuations. Extreme examples are cell-size variables and growth-rate, which exhibit super-diffusive scaling *γ* > 1/2 indicating persistent fluctuations, and correspondingly flexible set-points. In contrast *r* has a very small power of ≈ 0.3, suggesting it is anti-persistent, and correspondingly a tightly regulated set-point. These results indicate the different nature of each cellular variable, that is reflected by its various statistical properties on both long and short timescales.

## Discussion

The biological cell is a complex system with a vast number of microscopic degrees of freedom. At the mesoscopic scale, multiple phenotypic variables are dynamically coupled to one another. How gene expression, metabolism, growth, division, and other cell cycle events coordinate to give rise to growth homeostasis over extended time is a fundamental question; even for simple cells such as bacteria, much remains to be understood.

A central challenge when interpreting homeostasis in a multi-dimensional system is to identify which variables are regulated and which are constrained as a byproduct of interactions. Here, we addressed this question by developing a framework that decouples homeostatic set-points from temporal fluctuations around them. Given long enough single-cell measurements, the homeostatic set-points can be estimated by the temporal average of each variable. We propose that tightly regulated variables will have robust set-points that are buffered from environmental and other perturbations. Other, less regulated variables – while still possibly maintaining homeostasis in the sense of stable distributions – will exhibit a higher sensitivity to environmental changes, reflected by their set-points moving around with perturbations.

Our analysis relies on single-cell measurements from trapped *E. coli* in two types of microfluidic devices. Using the law of total variance and conditioning of data on individual traps, we separated the contribution of set-point variation from temporal fluctuations to the total variance. Examining a collection of phenotypic variables, we found a large range of behaviors, from highly flexible set-points that can move up to 30 – 40% of their values, to tightly controlled ones that vary at the level of sampling error or below (Fig. 2B). Making use of the recently developed sisters machine, we could assign fractions of the conditioned variance to lineage history separately from the effect of micro-environment. All variables showed a similar sensitivity to lineage history, about 2 – 5% of the total variance. This component can be mediated by noise in expression of metabolic enzymes; previous work has demonstrated that such noise can induce metabolic variability that manifests in growth-rate (Labhsetwar et al. 2013; Nikolic et al. 2017). However, this is only a minor contribution to the total variance, leaving the larger portion to the effect of trap environment. Thus, the graded conditioned variance of different cellular phenotypic variables reflects their wide range of sensitivities with respect to the environment.

The uniqueness of a microfluidic trap is defined in terms of several parameters, such as its temperature, medium composition, physical dimensions, and perfusion rate. While most studies routinely pool all data from experiments performed in the same controlled conditions, the limitations of repeatability across experiments was highlighted in recent studies (Yang et al. 2018; Kohram et al. 2021). In this study, we actually utilized the non-repeatability of trap environments to learn something qualitative about the cell itself. Thus, while we do not have control over the environmental perturbations, we can compare the behavior of different variables and how they co-vary. The qualitative picture arising from the variance decomposition is consistent across several data-sets that we have examined. In Fig.2 itself, panels B and C are measurements from two different microfluidic devices; SI Fig.8 shows the effect of pooling together different traps and different experiments, while SI Figs.10 and 9 apply our analysis to publicly available data from other labs, under nominally similar conditions. We find that the quantitative values of conditioned variance vary substantially between data-sets, but most qualitative observations made in this paper are robust. Future research might extend these comparisons and study different medium conditions and different bacterial species within the same framework.

Examining the correlations between the set-points of different variables, we found a pattern of correlation indicating a hierarchy of buffering. Time variables – growth-rate and generation time – have sloppy set-points that strongly co-vary with one another. Their product, logarithmic fold-growth, already buffers some of the external perturbations and exhibits a more robust set-point; it is itself negatively correlated with division fraction. Finally, their product defines the most protected variable – cell size ratio across consecutive cycles – which is almost perfectly buffered from all types of perturbations, at noise level or below. Based on this picture we predicted that set-points are confined to a manifold in the three-dimensional space (*α*, *τ*, *f*); this prediction was verified to high precision with a large set of data.

Our results suggest that, at the mesoscopic scale, phenotypic cellular variables exhibit a hierarchy of sloppiness. Moreover, we have identified combinations that become increasingly stiff and finally make up a control variable that is highly buffered from perturbations. The concept of sloppiness was first suggested in the context of parameter sensitivity of complex models (Daniels et al. 2008). Extending this notion beyond models, a sloppy dynamical system is characterized by a hierarchy of sensitivities for different variable combinations (Braun 2020). To our knowledge, this is the first explicit demonstration of such sloppy-system behavior of a biological cell from experimental data.

From a control-theory point of view, we have found that homeostasis does not have a single well-defined set-point but rather a “set manifold”, with different lineages maintaining homeostasis around points scattered inside the manifold. This suggests that some variables define null directions along which changes in time-averaged values do not alter the functionality of the system - namely, maintaining growth and division homeostasis. A similar concept was developed in the context of motor control-specifically the control of muscles by neural activity, where certain directions in the high-dimensional space of neural activity did not alter the muscle output and were therefore termed “output-null” directions (in contrast to “output-potent” ones, see (Kaufman et al. 2014)). In motor learning, the significance of such null-directions was implicated in acquiring novel capabilities while keeping functional structures intact (Perich, Gallego, and Miller 2018). Indication of a possible analogous role for our system is suggested by recent results on transitions between external conditions (Panlilio et al. 2020); this possibility calls for further investigation.

The fact that some lineage set-points in Fig. 4 are not at *f* = 1/2 but still on the homeostatic manifold, suggests that division fraction can participate in growth and division homeostasis. Our results show that it is strongly coupled to fold-growth such that the two are compensates, though we cannot distinguish which variables actively follows which. Recent results showed that, on the short-timescale of a single cell-cycle, growth rate compensates for the preceding fraction (Kohram et al. 2021); on the long-timescale of many generations, further investigation is needed to decipher the mechanisms involved. Biologically, a set-point for division which is consistently different from 1/2 might be an artefact of the trapping apparatus; the breaking of symmetry between daughter cell trajectories can reflect e.g. aging, since we follow the old-pole cell throughout the experiment; or a geometrical constraint, since the mother cell is confined at the bottom of the trap. Clearly, in a dividing population tree, following any particular lineage is arbitrary and division is expected to be symmetric along time. Nevertheless, our results demonstrate that division fraction has the capacity to contribute to homeostasis – or to be compensated for – when needed. They also highlight the gap that still remains between our understanding of a trapped lineage and of a population.

The high precision of the homeostatic manifold (Fig. 4) suggests that long-term growth and division homeostasis takes place without size control. However in the short-term, cell size clearly couples to growth within a cell-cycle. For example, recent work has shown that certain discrete events related to division depend on threshold accumulation of specific proteins, whose production is in turn dependent on cell size (Nordholt, Heerden, and Bruggeman 2020; Panlilio et al. 2020; Si et al. 2020). In other publications growth-rate was reported to compensate, within a single cycle, for noise that occurs at division (Vashistha, Kohram, and Salman 2021; Nordholt, Heerden, and Bruggeman 2020; Kohram et al. 2021). Such coupling induces correlations between cell size and effective per-cycle variables, e.g. division time and growth-rate; however, these could be an indirect consequence of basic biophysical size-dependent events rather than a regulation mechanism aimed at controlling cell size itself. In support of this view, we found that cell size is a sloppy variable that exhibits a flexible set-point, such that different lineages sustain homeostasis around different average cell sizes. Relative to the stiffest variable *r*, its set-point spans a range ~ 250 times larger. The short-timescale mechanisms involving cell size are possibly separate from the long-term ones that couple the homeostatic set-points to one another. Further work is required to test this hypothesis, as well as to decipher the underlying mechanisms on the long timescales.

Our work offers a general methodology for analyzing homeostasis in a multi-dimensional system from ensembles of temporal data, to reveal tightly controlled variables without prior assumptions. This approach can be widely used and is already being tested in other contexts. Applied to bacterial growth and division, our results taken together highlight the multiple-timescale and multi-variable nature of bacterial homeostasis. As more high-quality data that extends over long times accumulate, a future challenge is to test the generality of these observations to other cell populations that maintain stable growth and division.

## Methods

### Data Processing

Raw movies of dividing bacteria are analyzed to produce time-series of bacterial cell length inside microfluidic channels. To detect a division event from the length measurements, we calculated the difference between consecutive time-points and set a difference threshold that, if passed, means that a division event has most likely occurred. Having identified and verified these division events, we fit an exponential curve to the length measurements whose exponent is the growth-rate and whose intercept is the initial size of the cycle. Other phenotypic variables are derived accordingly (see Fig. 2A). The measurements are of high precision and low noise-level, and results are independent of precise definitions; for example, fold-growth can be defined alternatively as *ϕ* = *ατ* or ln (*x_τ_*/*x*_0_) with very similar results. Filamentation events were removed from the data by excluding cycles that contain variables 3 standard deviations away from the mean (using single-lineage statistics). Analysis was done in Python and the code is available at our repository on github: https://github.com/astawsky/Multiple-Timescales-in-Bacterial-Growth-Homeostasis.git.

### Artificial lineages

To estimate finite sampling effects, we create for each experiment a set of artificial lineages with the same statistics as the measured ones – number of traces and number of generations in each trace. The phenotypic variables are drawn at random from the pooled ensemble and therefore averaging over artificial lineages represents only the statistical effect of averaging over arbitrary finite groups of variables from the pool. This artificial data-set helps to assess the finite-size effects that emerge from the averaging procedure itself, without any temporal information in the actual lineages across time.

### Three-way Variance Decomposition

For pair lineages in a single sisters machine trap, the phenotypic variable *z* can be expressed as the sum of the trap average 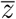, deviation of lineage-average from trap-average, and lineage-specific temporal fluctuations:

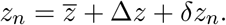

Where 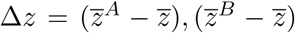 average to zero over the two neighboring lineages *A*, *B*, and *δz* averages to zero over time in each lineage, causing the cross-terms to vanish from the total variance:

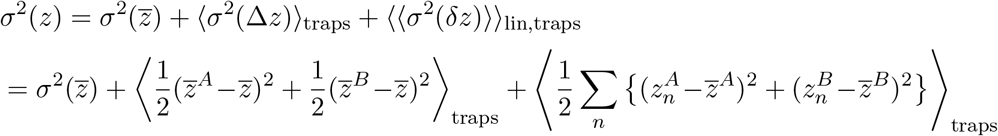

This is equivalent to Eq. 4, for the case of two lineages of equal lengths. In the more general case, averaging over lineages takes into account the number of cell-cycles in each one with the appropriate weight factor.

### Covariance Decomposition

Our variance decomposition for single trapped lineages, specifies how much of the pooled ensemble variance comes from long and short timescales:

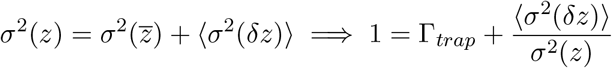

where *z* is phenotypic variable in a single trapped lineage, 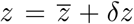. If we generalize this equation to the covariance between two phenotypic variables we get:

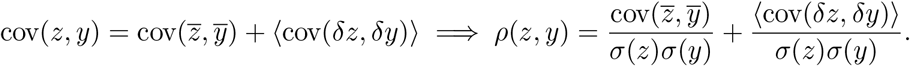

It is important to emphasize that neither component on the r.h.s.is itself a Pearson correlation, because they are normalized by the pooled standard deviations. In contrast, Pearson coefficients are normalized by their corresponding standard deviations:

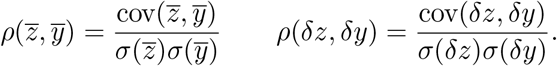

For example, we find cov(*α*, *τ*) = −.0083; with the standard deviations of the pooled ensemble *σ*(*α*) = 0.222*, σ*(*τ*) = 0.131 and the standard deviations of the time-averages themselves 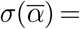 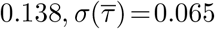, we find

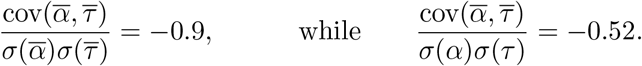

SI Fig. 11 presents the covariance decomposition for all pairs of phenotypic variables in the sisters machine data-set.

### Detrended Fluctuation Analysis (DFA)

We follow Peng et al. 1995 to characterize the accumulation of fluctuations in a time-series. For a window of a fixed number of cycles *k* inside the lineage containing *N* cycles let 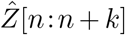 be the least-squares linear regression to the *Z*[*n*:*n* + *k*] points found inside each window. This line approximates the trend inside the window, and the root-mean-squared deviation of the data around the line is the detrended fluctuation:

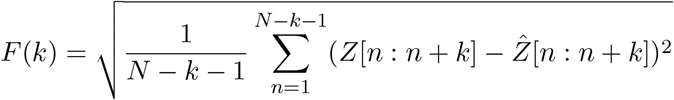

For 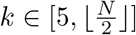, we fit via a least squares linear regression on the log-log graph of *F* (*k*) vs. *k*. In general *F* (*k*) ∝ *k^γ^*, where *γ* is the scaling exponent and tells us how the phenotypic variables of subsequent cycles are correlated. The window sizes increase in steps of 4 until the maximum window size and there is a 3 generation-gap between two windows of the same size. If a lineage has less than three windows sizes we can use to approximate the scaling exponent *γ*, we discard this lineage.

The need to use a detrended analysis to measure persistence comes from the fact that phenotypic variables in some lineages have a monotonic trend, and are therefore non-stationary (see SI Fig. 16). In this case, traditional spectral analyses such as the Hurst exponent, autocorrelation function and Fourier transform give slightly incorrect scaling because homeostasis is now around a linear trend, possibly imposed at least in part by a changing micro-environment. For lineages that do not have a trend, the results for DFA and the previously mentioned analyses are equivalent (Peng et al. 1995).

## Acknowledgements

This research was supported in part by the Israeli Science Foundation (N.B.) and Binational Science Foundation (joint N.B. and H.S.). We thank Omri Barak and Lee Susman for valuable comments on the manuscript. We thank Maryam Kohram for organizing the data from (Kohram et al. 2021).

## Supplementary Information

**Table 1:** Throughout this paper we refer to experiments and their conditioned ensembles by a unique ID shown in the first column. The second column shows the references of these experiments, which can be found in the bibliography. The third column shows the number of lineages in the experiment and the fourth column shows the mean and standard deviation of the statistics of the number of generations-per-lineage in each experiment. In all experiments, *E. coli* bacteria were grown in LB medium.

| Exp. | Reference | # of<br>lins. | # of<br>gens. | Temp.<br>(°C) | Strain | Date |
| --- | --- | --- | --- | --- | --- | --- |
| 0 | Vashistha 2021 | 48 | $20 \pm 13$ | 32 | MG1655 | 15/10/18 |
| 1 | Vashistha 2021 | 14 | $26 \pm 10$ | 32 | MG1655 | 27/06/18 |
| 2 | Vashistha 2021 | 10 | $18 \pm 4$ | 32 | MG1655 | 13/07/18 |
| 3 | Vashistha 2021 | 180 | $30 \pm 23$ | 32 | MG1655 | 28/07/18 |
| 4 | Vashistha 2021 | 178 | $22 \pm 11$ | 32 | MG1655 | 12/10/18 |
| 5 | Susman 2018 | 58 | $63 \pm 39$ | 32 | MG1655 | Pooled exps. |
| 6 | Korham 2020 | 30 | $58 \pm 10$ | 32 | MG1655 | Pooled exps. |
| 7 | Susman 2018 | 20 | $210 \pm 53$ | 32 | MG1655 | Pooled exps. |
| 8 | Tanouchi 2015 | 65 | $69 \pm 0$ | 25 | K-12 substr. MC4100 | |
| 9 | Tanouchi 2015 | 54 | $71 \pm 2$ | 27 | K-12 substr. MC4100 | |
| 10 | Tanouchi 2015 | 160 | $69 \pm 0$ | 37 | K-12 substr. MC4100 | |
| 11 | Wang 2010 | 31 | $107 \pm 48$ | 37 | MG1655 (CGSC 6300) | 10/02/09 |
| 12 | Wang 2010 | 126 | $98 \pm 35$ | 37 | MG1655 (CGSC 6300) | 29/01/09 |
| 13 | Wang 2010 | 174 | $142 \pm 51$ | 37 | MG1655 (CGSC 6300) | 02/07/09 |
| 14 | Wang 2010 | 56 | $90 \pm 56$ | 37 | MG1655 (CGSC 6300) | 31/01/09 |
| 15 | Wang 2010 | 315 | $96 \pm 54$ | 37 | MG1655 (CGSC 6300) | 25/05/09 |
| 16 | Wang 2010 | 228 | $67 \pm 41$ | 37 | MG1655 (CGSC 6300) | 12/05/09 |
| 17 | Wang 2010 | 141 | $32 \pm 26$ | 37 | MG1655 *lexA3* | 30/09/09 |
| 18 | Wang 2010 | 185 | $38 \pm 36$ | 37 | MG1655 *lexA3* | 23/09/09 |
| 19 | Wang 2010 | 164 | $35 \pm 23$ | 37 | MG1655 *lexA3* | 22/09/09 |

**Table 2:** The Coefficient of Variation, defined as the standard deviation over mean, for the phenotypic variables over all bacterial cell cycles, measured in the different experiments (corresponding to the experiments described in SI Table 1). Each entry lists the CV over the pooled ensemble of each experiment. Bottom line lists the average and STD over all experiments.

| Exp. | $r$ | $\phi$ | $f$ | $\Delta$ | $\tau$ | $x_0$ | $x_\tau$ | $\alpha$ |
| --- | --- | --- | --- | --- | --- | --- | --- | --- |
| 0 | 0.12 | 0.19 | 0.06 | 0.20 | 0.22 | 0.15 | 0.13 | 0.11 |
| 1 | 0.12 | 0.19 | 0.07 | 0.22 | 0.21 | 0.13 | 0.12 | 0.13 |
| 2 | 0.21 | 0.30 | 0.14 | 0.33 | 0.37 | 0.19 | 0.18 | 0.30 |
| 3 | 0.15 | 0.23 | 0.08 | 0.26 | 0.24 | 0.18 | 0.17 | 0.15 |
| 4 | 0.14 | 0.20 | 0.06 | 0.23 | 0.22 | 0.17 | 0.16 | 0.13 |
| 5 | 0.18 | 0.30 | 0.14 | 0.34 | 0.34 | 0.23 | 0.21 | 0.24 |
| 6 | 0.13 | 0.21 | 0.07 | 0.24 | 0.23 | 0.16 | 0.15 | 0.15 |
| 7 | 0.16 | 0.24 | 0.08 | 0.27 | 0.27 | 0.17 | 0.16 | 0.14 |
| 8 | 0.18 | 0.25 | 0.08 | 0.30 | 0.27 | 0.25 | 0.22 | 0.08 |
| 9 | 0.16 | 0.20 | 0.08 | 0.24 | 0.20 | 0.15 | 0.15 | 0.18 |
| 10 | 0.11 | 0.16 | 0.06 | 0.17 | 0.17 | 0.14 | 0.12 | 0.07 |
| 11 | 0.16 | 0.23 | 0.06 | 0.31 | 0.25 | 0.21 | 0.22 | 0.08 |
| 12 | 0.14 | 0.21 | 0.05 | 0.28 | 0.23 | 0.22 | 0.21 | 0.08 |
| 13 | 0.17 | 0.27 | 0.07 | 0.35 | 0.29 | 0.25 | 0.24 | 0.11 |
| 14 | 0.17 | 0.25 | 0.07 | 0.34 | 0.28 | 0.26 | 0.26 | 0.13 |
| 15 | 0.16 | 0.24 | 0.08 | 0.34 | 0.26 | 0.25 | 0.25 | 0.10 |
| 16 | 0.27 | 0.35 | 0.13 | 0.50 | 0.37 | 0.30 | 0.34 | 0.11 |
| 17 | 0.15 | 0.24 | 0.05 | 0.29 | 0.27 | 0.17 | 0.16 | 0.11 |
| 18 | 0.14 | 0.23 | 0.05 | 0.28 | 0.28 | 0.19 | 0.18 | 0.16 |
| 19 | 0.15 | 0.26 | 0.05 | 0.30 | 0.26 | 0.18 | 0.18 | 0.11 |
| CV | 0.16 ±<br>0.04 | 0.24 ±<br>0.04 | 0.08 ±<br>0.03 | 0.29 ±<br>0.07 | 0.26 ±<br>0.05 | 0.2 ±<br>0.05 | 0.19 ±<br>0.05 | 0.13 ±<br>0.06 |

**Figure 8:**
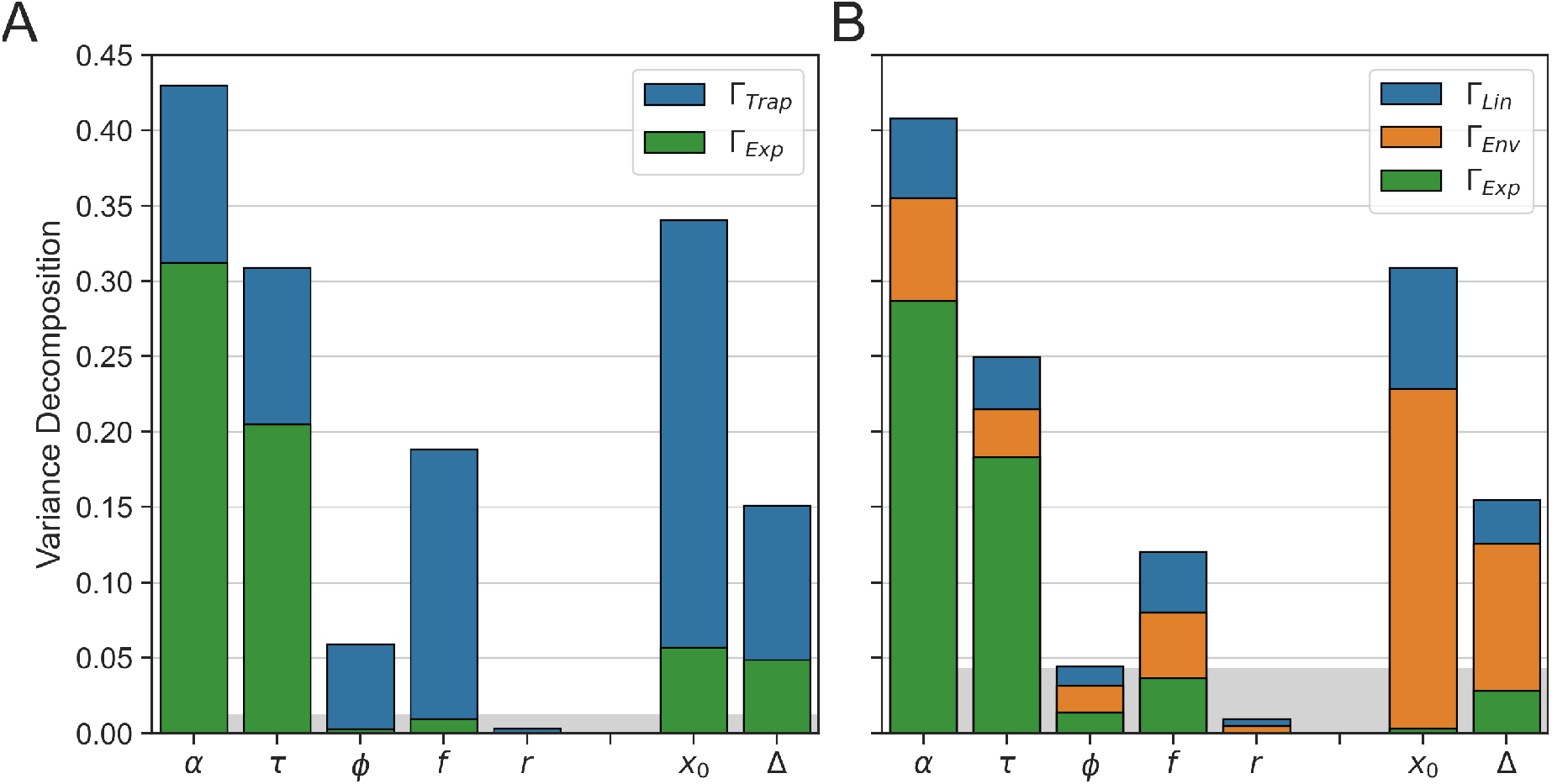
Variance decomposition effect of experiments. The same data as in Fig. 2B,C but using the law of total variance to decompose how much of the total variance can be attributed to different experiments. This is done by an extension of the conditioning which includes also individual experiments. If different experiments are clustered in phenotypic phase-space then the green bar should be more than half the height of the blue bar. The opposite would be true if experiments are not clustered. In **(A)** we can see that there is strong clustering per experiment for growth-rate and inter-division time and lower clustering per experiment for initial size and added length. In **(B)** we see that the majority of the clustering for initial size and added length comes from the trap variability in each experiment separately.

**Figure 9:**
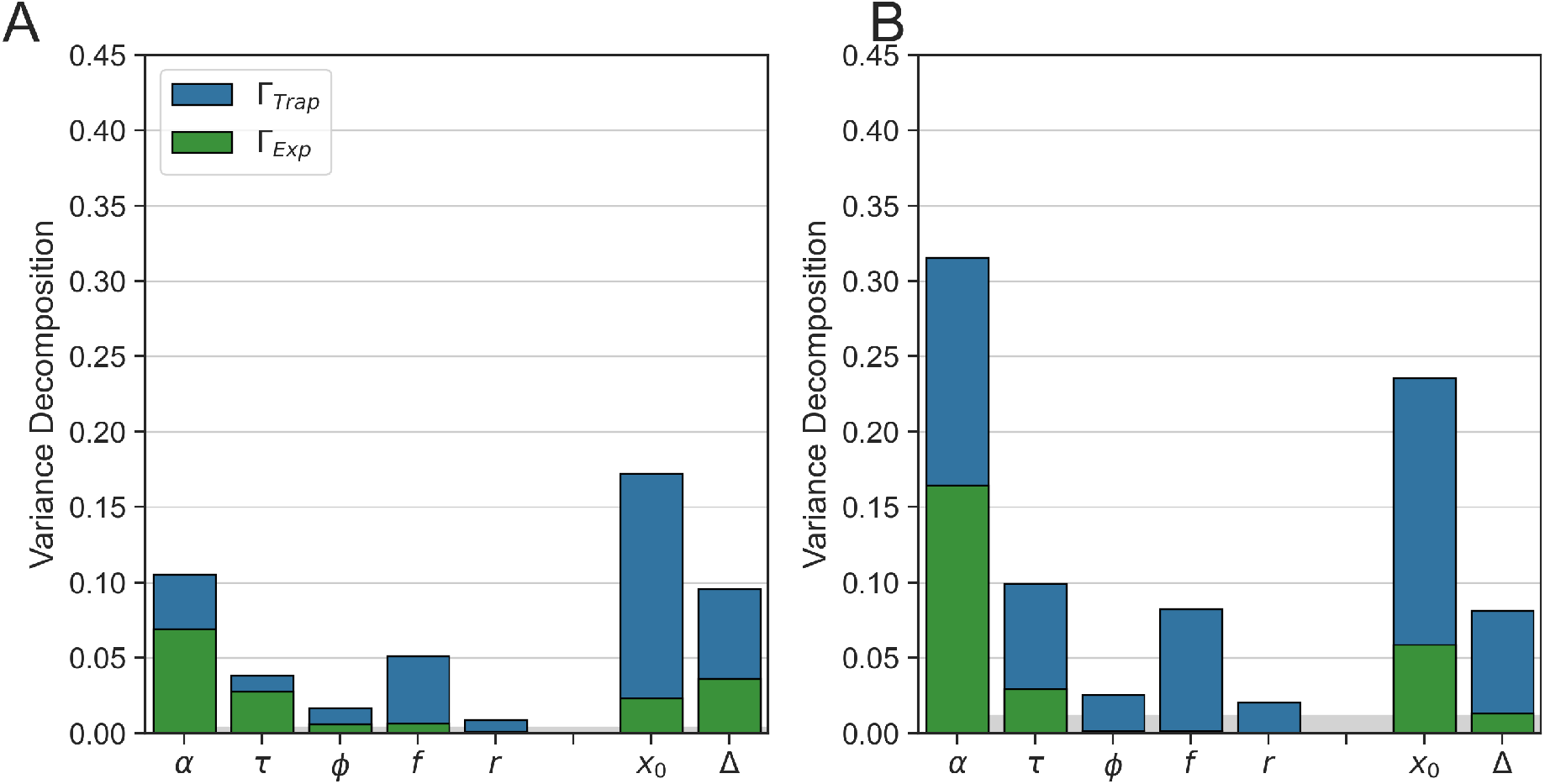
Variance decomposition across experiments for two bacterial strains. Same analysis as in SI Fig. 8A, applied to mother machine data from experiments 11 – 16 and 17 – 19 respectively, see SI Table 1.

**Figure 10:**
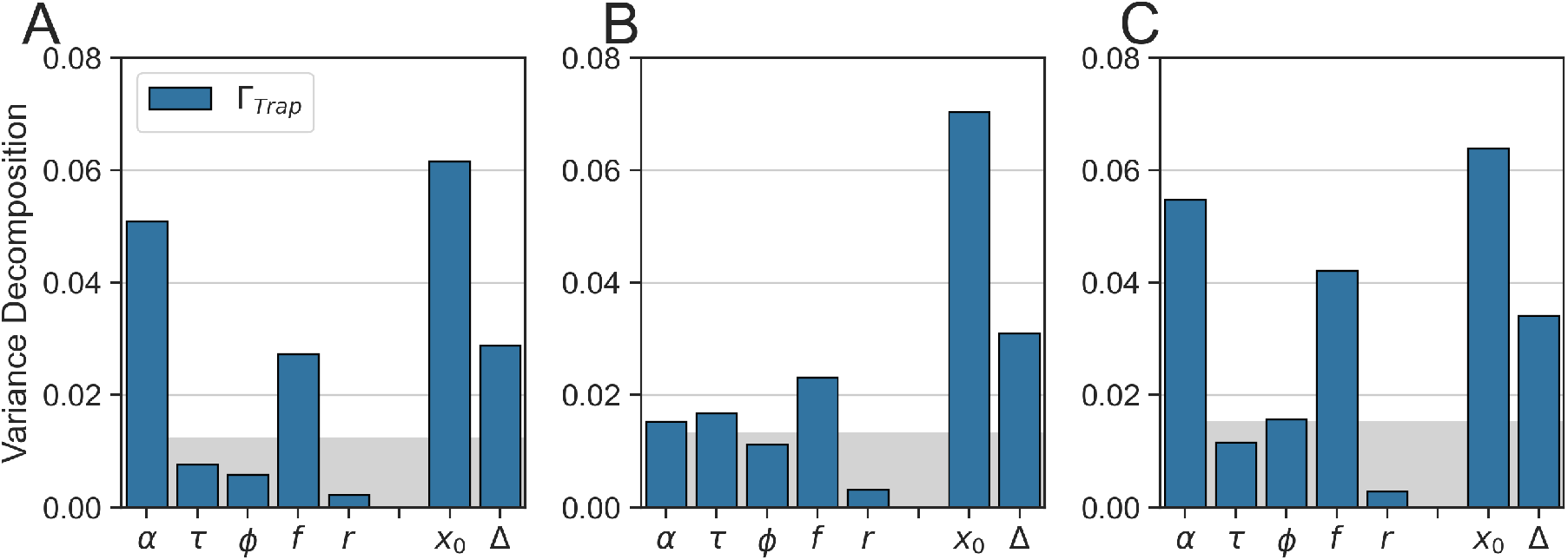
Variance decomposition at three temperatures. This figure shows results of the same analysis as Fig. 2B, performed on mother machine experiments from Tanouchi et al. 2015 (experiments 8-10 in SI Table 1). Bacteria were grown in 25, 27 and 37°*C* in **A,B,C** respectively. All values of Γ_*trap*_ in this set of experiments are particularly small compared to other experiments examined (See Fig. 2 and SI Fig. 9 below). The relative values share some of the qualitative properties seen in the other data-sets.

**Figure 11:**
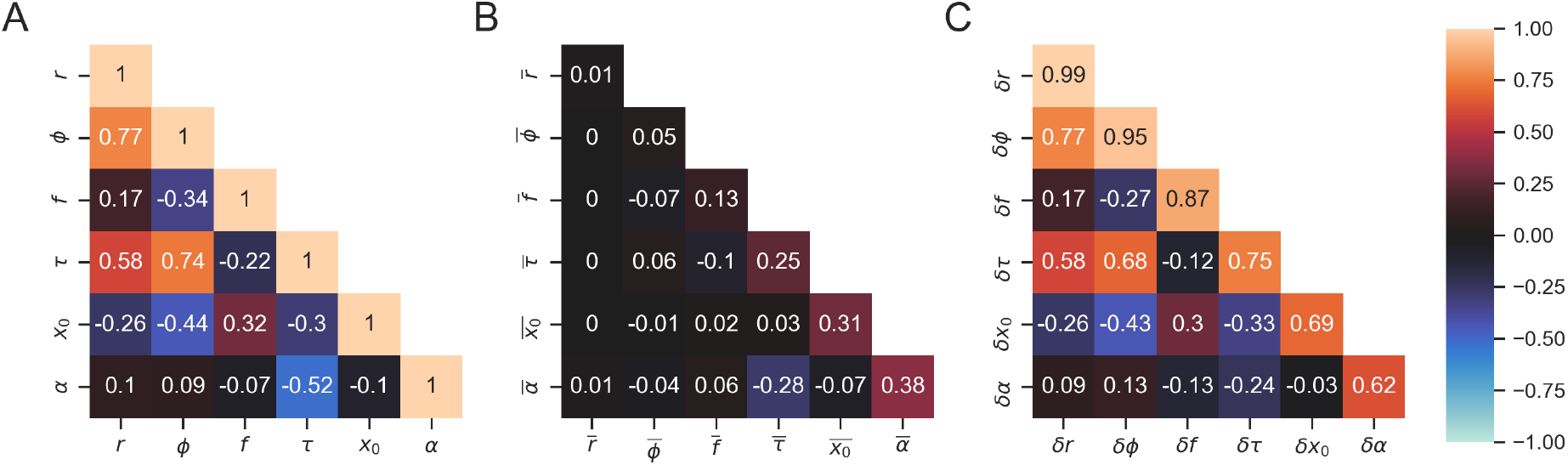
Covariance decomposition in individual lineages. **(A)** Normalized covariance (Pearson correlation coefficient) for all pairs of phenotypic variables over the pooled ensemble. **(B)**,**(C)** are the decomposition of entries in **A** to long-term and short-term contributions, respectively (see Methods). Entries on the diagonal are exactly the variance decomposition presented in Fig. 2, showing their hierarchy of conditioned variance from the stiffest variable *r* to the most sloppy one *α*. At the non-diagonal elements, the entry in **A** is the sum of entries in **B**,**C**, as expected. Examining the relative contributions of the two timescales, we find that *α* and *τ* have a significant long timescale contribution; other pairs of variables co-vary mostly on the short timescale contributions. In particular, the correlations of cell size with all other variables comes primarily from short-time, relative per-cycle variables.

**Figure 12:**
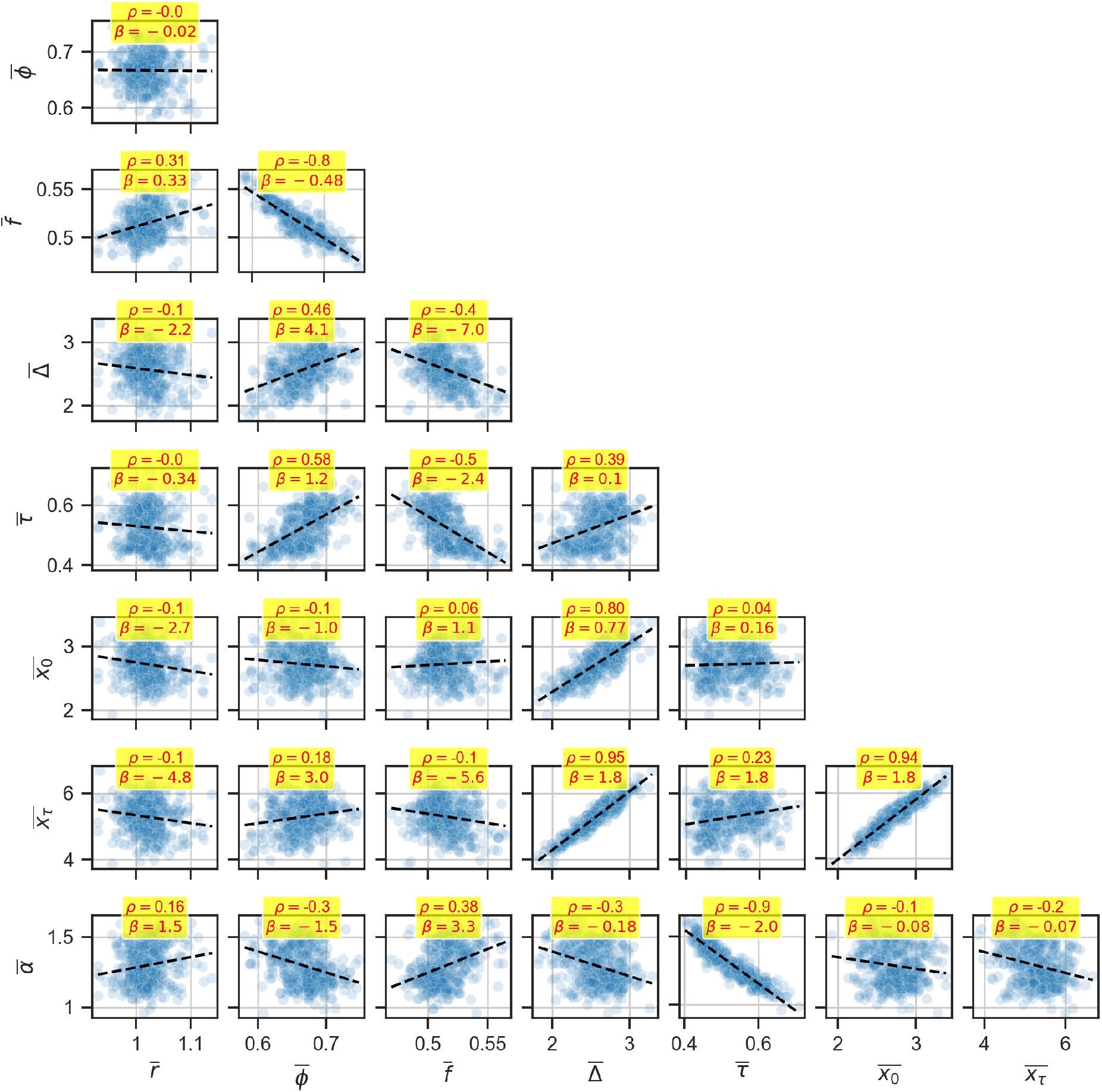
Long timescale correlations. This matrix of scatterplots shows the Pearson correlation (*ρ*) and best-fit slope (*β*) between a lineage’s time-average of two different phenotypic variables shown in the x- and y-axis labels. Every point represents a lineage longer than 15 generations found in the pooled ensemble of experiments 0-4, i.e. Sisters Machine experiments. There are 428 points. The strongest correlations seen here are 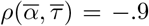, 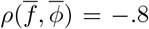, 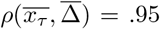, 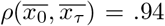, and 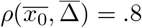. Also notice that 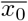 is not correlated with either 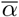 or 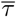, suggesting a regulation mechanism that does not measure size.

**Figure 13:**
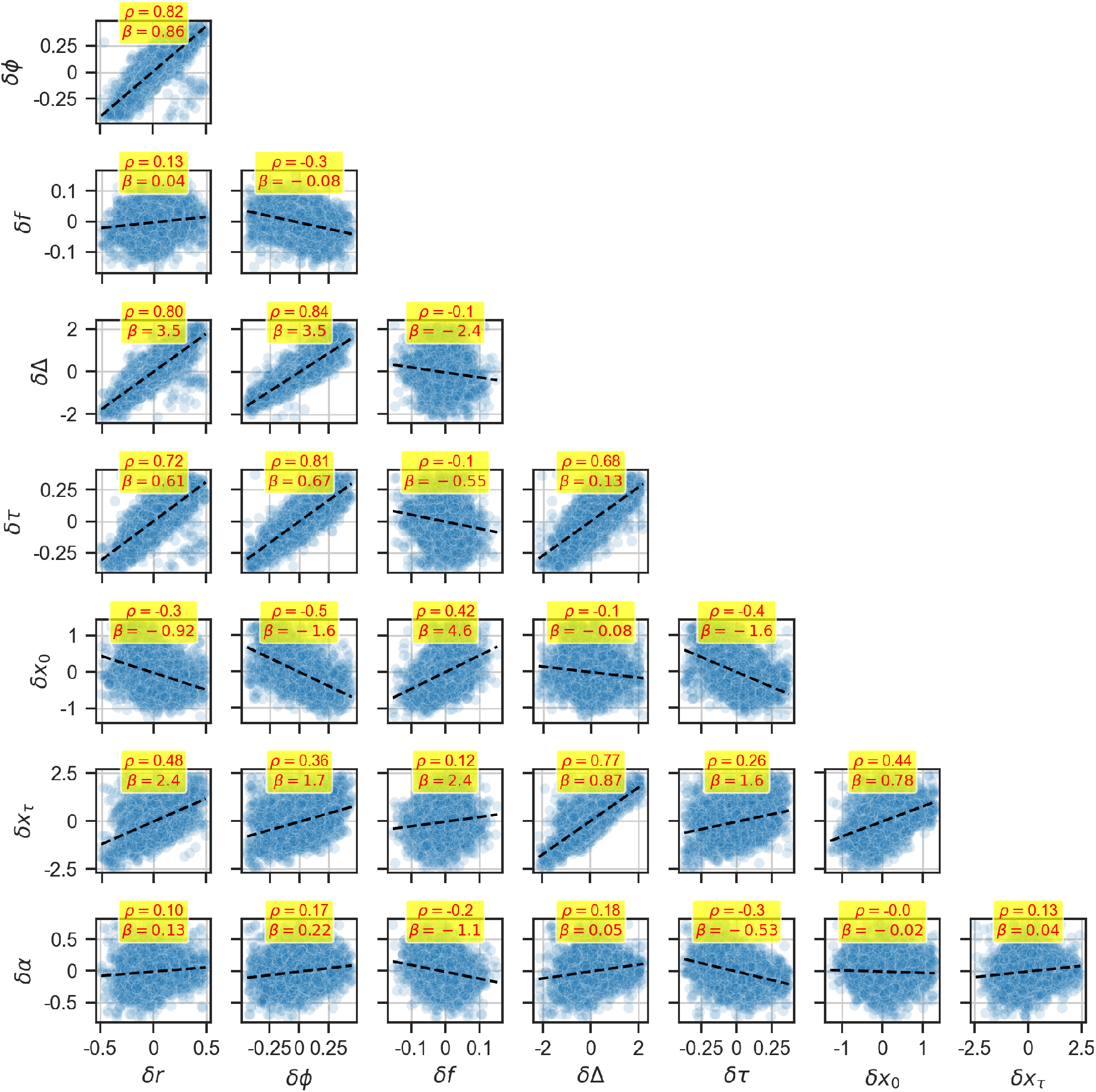
Short timescale correlations. This matrix of scatter-plots shows the Pearson correlation (*ρ*) and best-fit slope (*β*) between lineage-centered values of different pairs of phenotypic variables shown in the x- and y-axis labels. Every point represents a single cell cycles measurement in the pooled ensemble of experiments 0-4 (sisters machine experiments). There are a total of 10,900 points.

**Figure 14:**
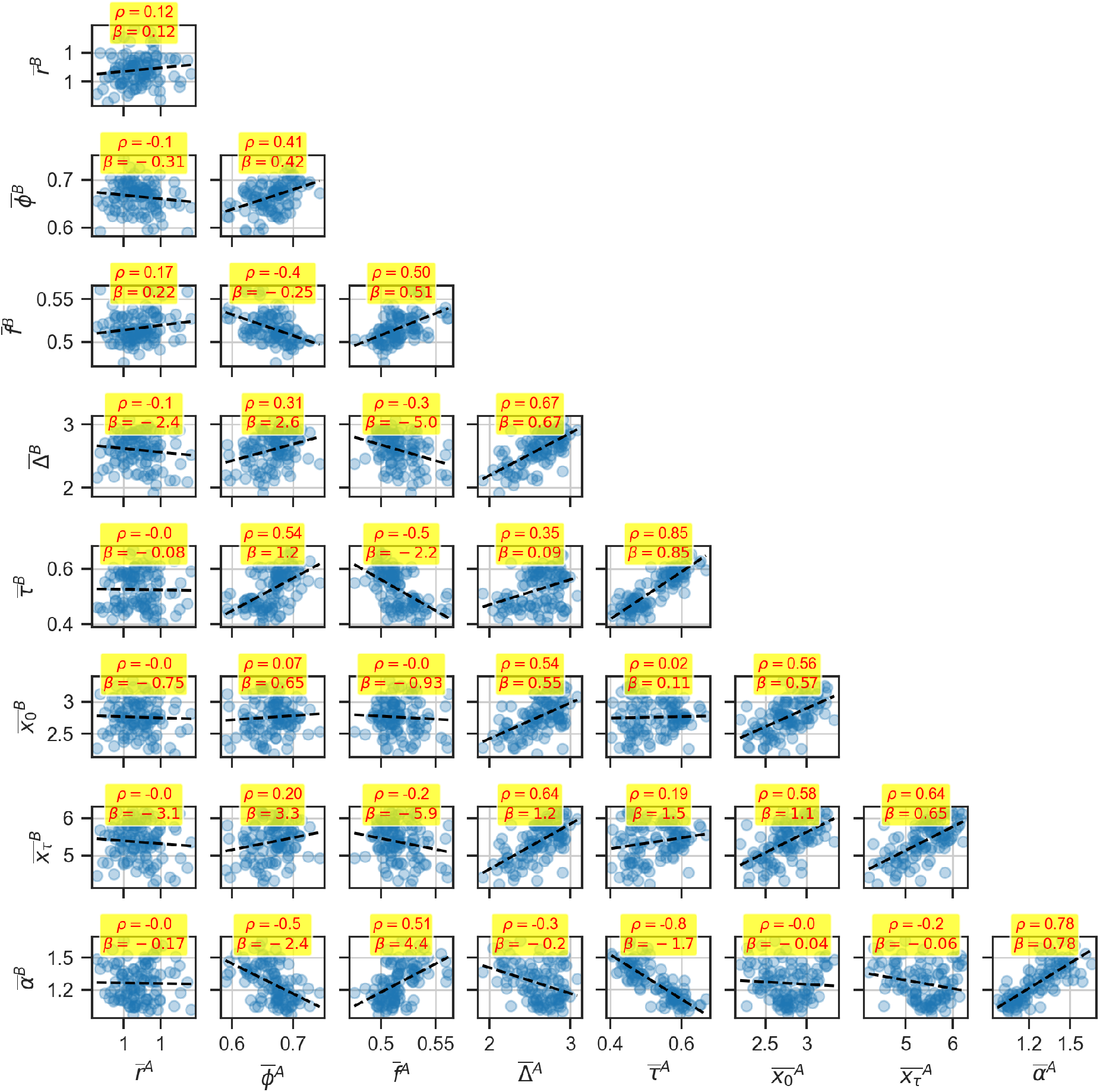
Long timescale correlations between neighbor lineages. This matrix of scatter-plots shows the Pearson correlation (*ρ*) and best-fit slope (*β*) between a neighbor lineage set-points of phenotypic variables denoted in the x- and y-axis labels. Every point represents a lineage longer than 15 generations from the sisters machine experiments. There are between 106 108 points in each panel. All variables except for *r* experience some micro-environmental influence because of the strong correlations found in the diagonal entries. In the off-diagonal, some correlations that are important to highlight are 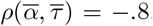, 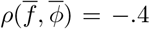, 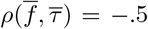 and 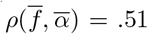. Moreover, it is important to note that 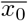 only correlates with itself and other size variables, of which it is a factor. This lack of correlation between size and growth variables suggests that micro-environment affects the two independently on the long-timescale.

**Figure 15:**
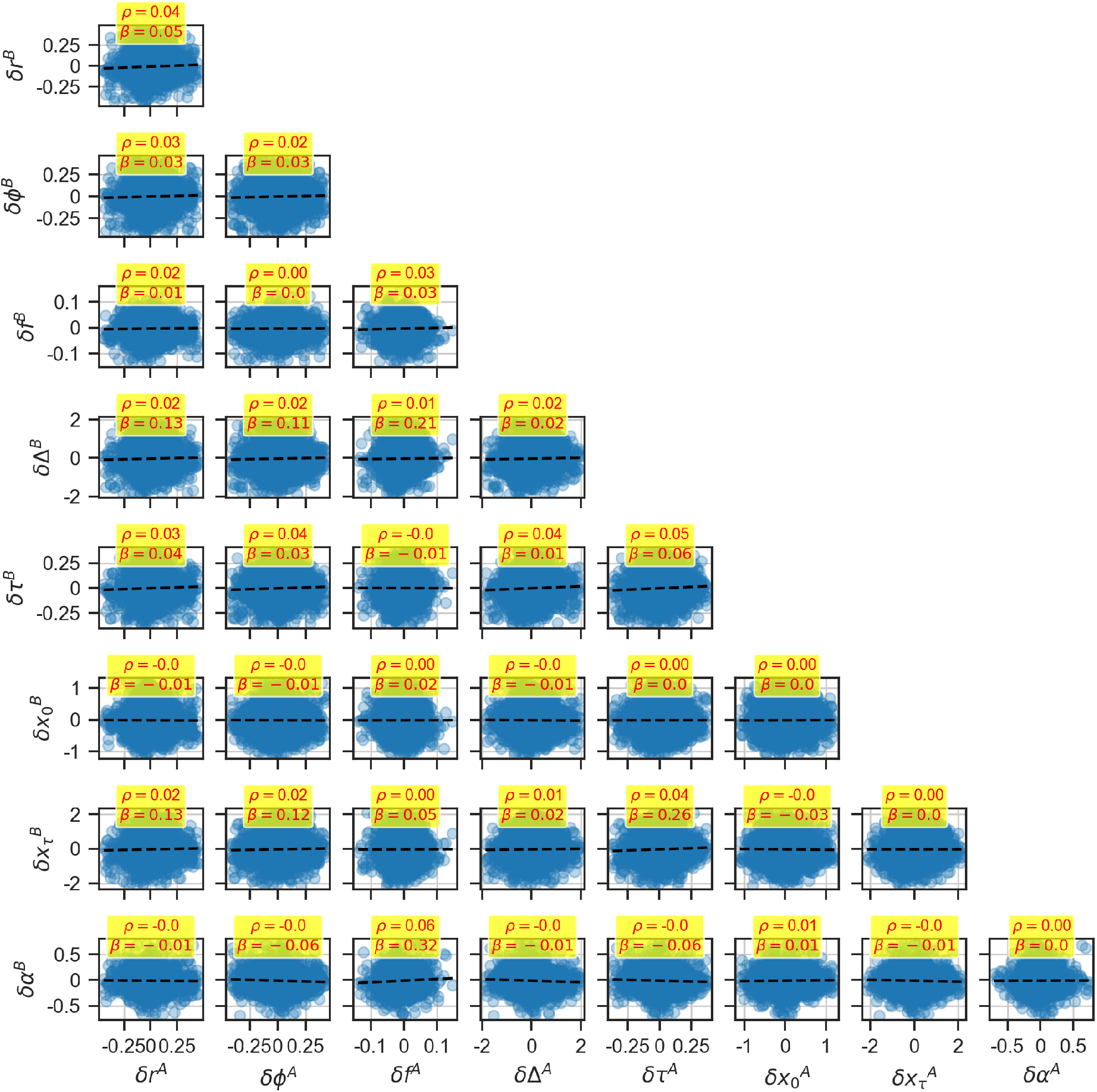
Short timescale correlations between neighbor lineages. This matrix of scatter-plots shows the Pearson correlation (*ρ*) and best-fit slope (*β*) between lineage-centered values of neighbor lineages for phenotypic variable pairs corresponding to the x- and y-axis labels. Every point represents a single bacteria in the pooled ensemble of experiments 0-4, ie. Sisters Machine experiments. There are 10,900 points. Every point represents neighbor bacteria of the same generation in the pooled ensemble of the sisters machine experiments. There are between 3430 − 3667 points in each panel. It is clear to see that there are no short timescale correlations between neighbor pair bacteria. This means that the micro-environment only influences the set-points of a lineage and not the fluctuations around it.

**Figure 16:**
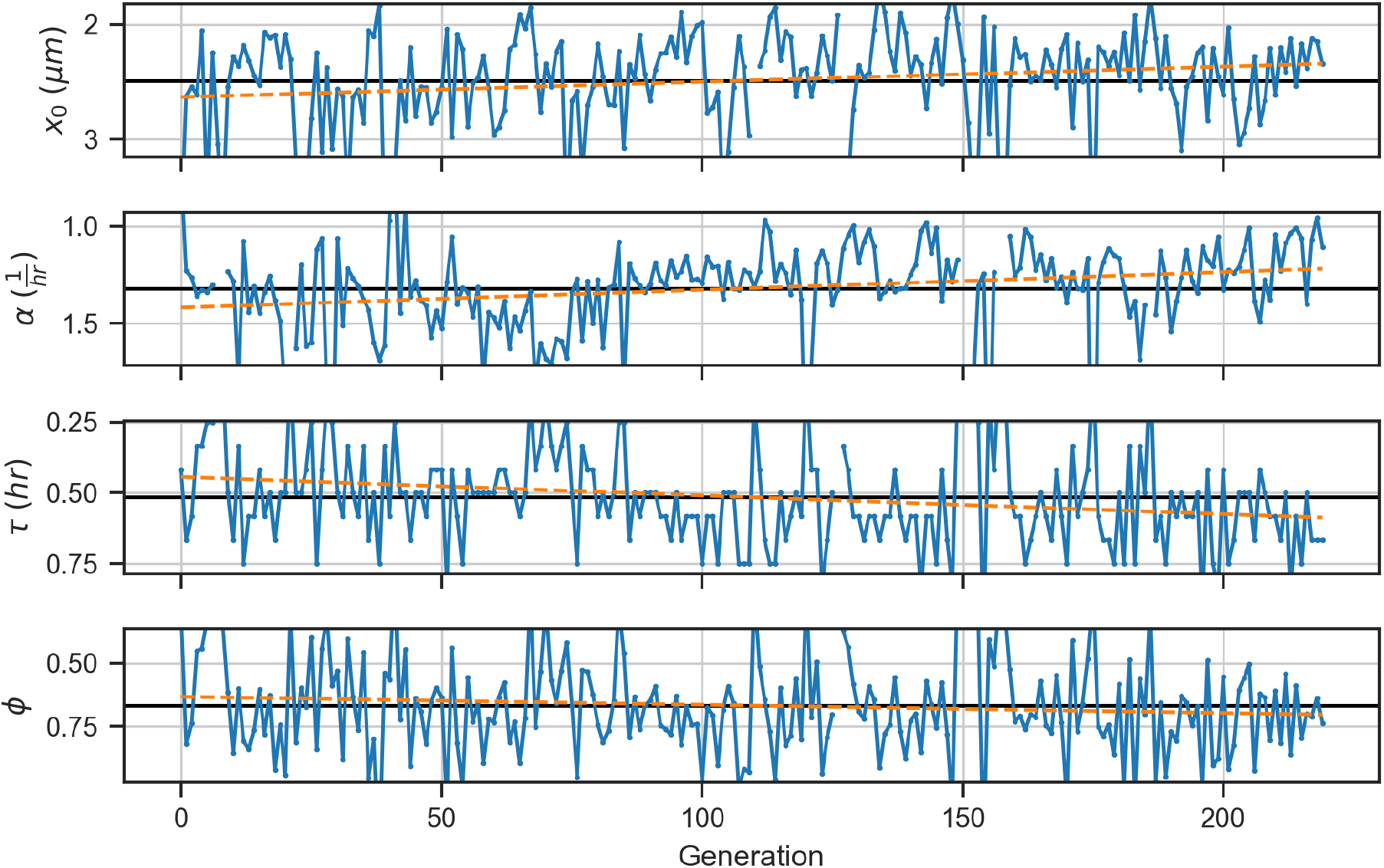
Trend of phenotypic variables. For a bacterial lineage of 207 generations found in experiment 7 (see Table 1), we can see a strong trend in both initial size and growth-rate across generations. The black line is the average across all generations and the orange line is the best fit line across time. Notice how the opposing trends for *α* and *τ* cancel out the trend in *ϕ*, consistent with our results of long timescale correlations.

**Figure 17:**
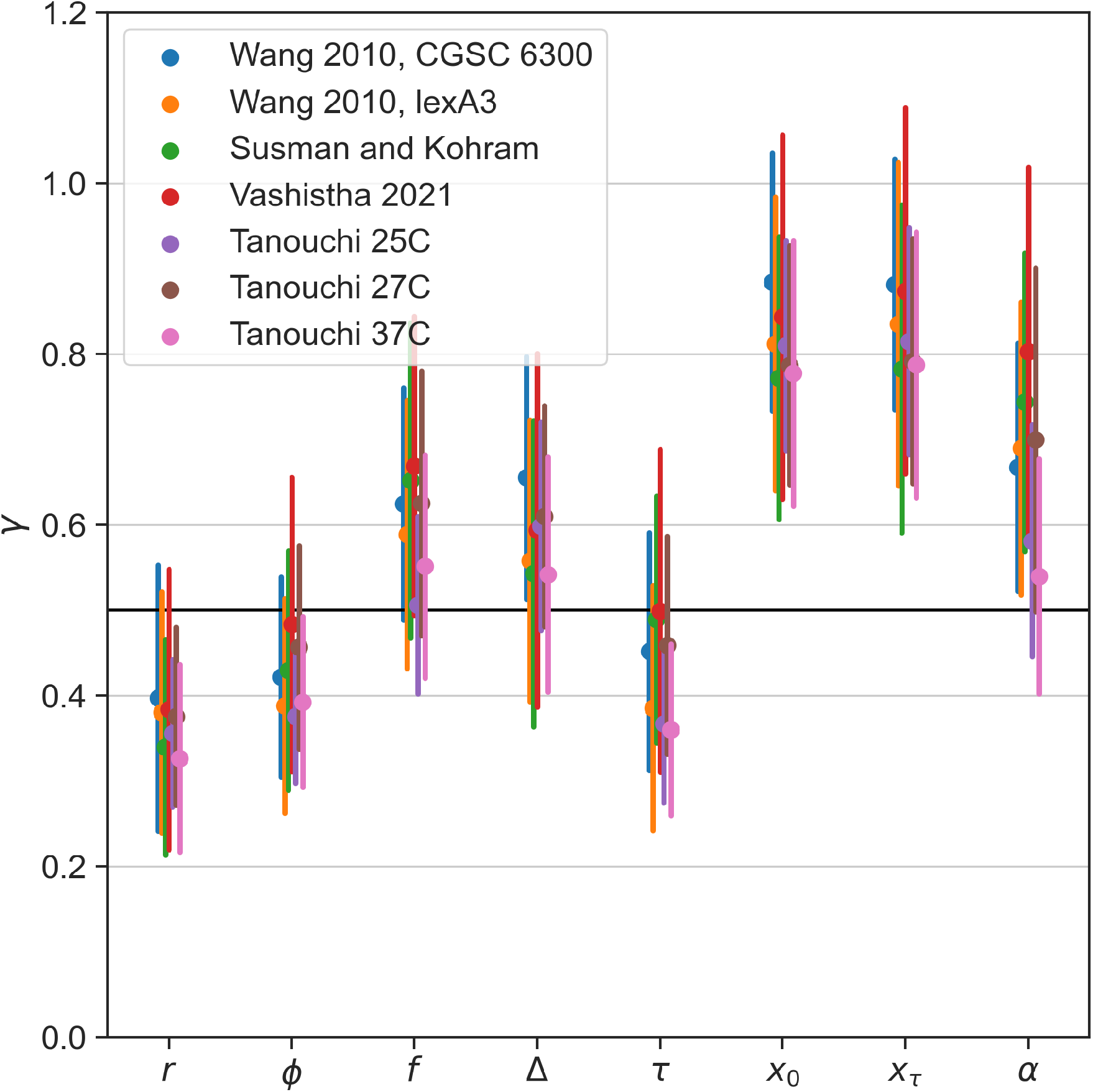
Persistence of phenotypic variables. This figure is related to Fig.7C but instead of pooling experiments, it shows the trace lineage scaling exponents for different experiment groups independently. While there is still some variance, the experiment-specific results are consistent with Fig.7C.

## Notes

https://github.com/astawsky/Multiple-Timescales-in-Bacterial-Growth-Homeostasis.git

### Competing Interest Statement

The authors have declared no competing interest.

